# Using a large-scale biodiversity monitoring dataset to test the effectiveness of protected areas at conserving North-American breeding birds

**DOI:** 10.1101/433037

**Authors:** Victor Cazalis, Soumaya Belghali, Ana S.L. Rodrigues

## Abstract

Protected areas currently cover about 15% of the global land area, and constitute one of the main tools in biodiversity conservation. Quantifying their effectiveness at protecting species from local decline or extinction involves comparing protected with counterfactual unprotected sites representing “what would have happened to protected sites had they not been protected”. Most studies are based on pairwise comparisons, using neighbour sites to protected areas as counterfactuals, but this choice is often subjective and may be prone to biases. An alternative is to use large-scale biodiversity monitoring datasets, whereby the effect of protected areas is analysed statistically by controlling for landscape differences between protected and unprotected sites, allowing a more targeted and clearly defined measure of the protected areas effect. Here we use the North American Breeding Bird Survey dataset as a case study to investigate the effectiveness of protected areas at conserving bird assemblages. We analysed the effect of protected areas on species richness, on assemblage-level abundance, and on the abundance of individual species by modelling how these metrics relate to the proportion of each site that is protected, while controlling for local habitat, altitude, productivity and for spatial autocorrelation. At the assemblage level, we found almost no relationship between protection and species richness or overall abundance. At the species level, we found that forest species are present in significantly higher abundances within protected forest sites, compared with unprotected forests, with the opposite effect for species that favour open habitats. Hence, even though protected forest assemblages are not richer than those of unprotected forests, they are more typical of this habitat. We also found some evidence that species that avoid human activities tend to be favoured by protection, but found no such effect for regionally declining species. Our results highlight the complexity of assessing protected areas effectiveness, and the necessity of clearly defining the metrics of effectiveness and the controls used in such assessments.

## Introduction

The increasing human footprint on natural ecosystems is leading to major declines in species’ populations (McRae et al., 2016) and has already resulted in thousands of extinctions (IUCN, 2018), to such an extent that Ceballos et al. (2017) characterised current times as a period of “biodiversity annihilation”. Habitat loss and degradation are the most important pressures on biodiversity (Vié et al., 2009; Balmford and Bond, 2005), as a result of anthropogenic activities such as agriculture, urbanisation, industry, transport and recreation (Foley et al., 2005). The most evident response to these threats is to establish areas with restricted, or even no human activities, *i.e.*, to create protected areas (PAs). Modern PAs have their origins in the 19^th^ century and currently represent the most important conservation tool, with about 15% of the global land area already protected to some extent, and coverage planned to reach 17% by 2020 (UNEP-WCMC IUCN, 2016).

Understanding the extent to which PAs are effective as biodiversity conservation tools is thus fundamental for guiding future conservation efforts. Accordingly, there is a substantial literature on PA effectiveness: as of the 1^st^ October 2018, 260 publications in the Web of Science included in their title “protected AND area* AND effective*”. However, within this literature there are disparate approaches to the concept of “effectiveness”.

A first set of studies questions whether PAs are effective at representing species or ecosystems, using gap analyses for measuring the overlap between PAs and the distributions of species or of ecosystem types (e.g., Rodrigues et al., 2004; Brooks et al., 2004). These studies do not directly quantify the effectiveness of PAs at conserving biodiversity, but the extent to which species or ecosystems are buffered from human impacts under the assumption that PAs are highly effective in doing so. A second set of studies focuses on the means employed locally by PA managers in order to protect biodiversity, for example in terms of staff or money (*e.g.,* Leverington et al., 2010). These analyses do not directly measure PA effectiveness in reducing human impacts, but rather the resources allocated to this purpose. A third type of studies quantifies the effectiveness of PAs at preventing the conversion of natural ecosystems, typically by comparing land use change (e.g., deforestation rates) in protected versus unprotected areas (e.g., Nelson and Chomitz, 2009; Andam et al., 2008). These studies quantify PA effects at the habitat or ecosystem level, rather than at the species level. Finally, a set of analyses focuses on measuring the effect of PAs on species themselves, either on the diversity of assemblages or on the abundance of individual species, typically by contrasting protected versus unprotected sites. This fourth approach to PA effectiveness is the focus of the present study.

The effectiveness of PAs at conserving species can be assessed by comparing population trends inside and outside PAs (e.g. Gamero et al., 2017; Devictor et al., 2007; Pellissier et al., 2013). Indeed, if PAs are effective, populations inside these areas are expected to be better buffered from threats and thus to decline less, or even to increase more, than those outside. Trends however can be misleading, because they are calculated in relation to a reference date (that seldom precedes all anthropogenic impacts) and because they are measured as percentages (which emphasise changes in small numbers). In this study, we focus instead on measures of PA effectiveness that assess current state, namely by contrasting population abundances and species diversity inside versus outside PAs (e.g. Coetzee et al., 2014; Kerbiriou et al., 2018; Devictor et al., 2007). These measures combine two types of effects: the effectiveness at selecting as PAs sites of higher-than-average conservation interest (i.e. differences that existed at the time of PA designation); and effectiveness at maintaining species richness and abundance within existing PAs (i.e. differences established subsequently to PA designation).

Three recent meta-analyses investigated the effects of PAs on the state of species abundance and/or diversity by synthesising studies that made pairwise comparisons between protected and unprotected sites (Geldmann et al., 2013; Coetzee et al., 2014; Gray et al., 2016). The underlying studies used in these meta-analyses did not necessarily aim to measure PA effectiveness; more often they investigated the effects of anthropogenic pressure, using PAs as benchmarks (e.g. Sinclair et al., 2002; Bihn et al., 2008; Wunderle et al., 2006, all integrated in Coetzee meta-analysis). In the meta-analyses, unprotected sites were treated as counterfactuals to the protected sites (i.e., by assuming that the latter would be in a similar condition to the former if it had not been protected), measuring the effect of protection as the observed difference between the two. These pairwise comparisons often contrast neighbouring sites, which presents the advantage of ensuring that both have broadly similar environmental characteristics (e.g. same climate). However, they do not necessarily take into account the fact that PAs tend to be biased in their location towards higher altitudes and lower productivity areas (Joppa and Pfaff, 2009). To reduce these biases, Gray et al. (2016) controlled for the differences in altitude, slope and agricultural suitability. Controlling for these factors means that their results are less influenced by PAs’ location biases and, therefore, that they reflect more strongly the effects of protection itself. Another potential confounding effect in pairwise comparisons of neighbouring sites arises from the leakage effect, whereby the human activities that would have taken place inside a PA are displaced to areas around it, artificially inflating the perceived effectiveness of PAs (Ewers and Rodrigues, 2008). This effect is difficult to control for, but should be reduced if the counterfactual sites are not immediately adjacent to the PAs.

An important decision when choosing a suitable spatial counterfactual to a PA, one that strongly affects the definition and thus the measure of PA effectiveness, is whether to control for habitat type or not. On the one hand, not controlling for habitat can lead to comparing sites that are not expected to have similar biodiversity regardless of their protection status (e.g. protected grasslands vs unprotected forests). On the other hand, controlling for habitat type can result in an overlooking of the effects that PAs have on biodiversity by preventing habitat changes (e.g. deforestation or urbanisation). For instance, given a hypothetical PA covering a natural grassland, possible counterfactuals include an unprotected site of similar habitat (i.e., natural grassland), an unprotected site with a different type of natural habitat (e.g., forest, wetland), or an unprotected site with human-modified habitat (e.g., extensive pasture, herbaceous cropland, urban area). The choice of counterfactual is certain to have a major impact on the differences observed, and thus on the measure of PA effectiveness, but it is not necessarily obvious which option is the best counterfactual. In theory, it is the site that best represents “what would have happened to the PA in the absence of protection”; in practice, this is not necessarily easily determined. All three metaanalyses (Geldmann et al., 2013; Coetzee et al., 2014; Gray et al., 2016) include comparisons where habitat has not been controlled for, meaning that the counterfactual’s habitat may be different or similar to the protected site’s habitat. Additionally, a subset of Gray et al. (2016)’s analyses focuses on comparisons between protected and unprotected sites with matched habitats. In the latter, the measure of PA effectiveness concerns protection from habitat degradation rather than protection from habitat conversion.

Another key consideration in analysing PA effectiveness is the biodiversity metrics applied to comparing protected and unprotected sites. The three meta-analyses employed a diversity of metrics, some at the level of species’ assemblages, some focused on individual species. Gray et al. (2016) used only assemblage-level metrics and found higher species richness and overall abundance inside PAs than outside, but no difference in rarefaction-based richness (i.e. number of species for a given number of individuals) nor in the proportion of endemic species. When matching sites with similar habitats, species richness was only higher in young and small PAs than in unprotected sites (no difference between other protected and unprotected sites), suggesting that the effect of PAs in preventing habitat degradation was light. Conversely, Geldmann et al. (2013) considered only species-level metrics (presence, abundance, nest survival) and found contrasted but mainly positive effects of PAs. Finally, Coetzee et al. (2014) considered both levels; at the assemblage level, they found higher species richness and overall abundance in protected than in unprotected sites; at the species level, they found that individual species abundances were typically higher inside PAs.

An alternative to measuring PA effectiveness through pairwise comparisons is to use statistical models in which covariates control for differences between protected and unprotected sites. This approach requires access to a large dataset on the spatial distribution of biodiversity, but reduces the subjectivity in the choice of counterfactuals, by making explicit which variables are controlled for, and the measure of effectiveness being investigated. For example, Devictor et al. (2007) applied this approach to survey data on common birds across France to find a positive effect of PAs on bird abundances for half of the species investigated, especially declining species. Duckworth and Altwegg (2018), working on bird abundance data collected across South Africa, found that PA coverage was positively correlated with occupancy of frugivorous, insectivorous, vegivores and predator species, and negatively correlated with occupancy of granivorous species.

In the present study, we quantify the effectiveness of Protected Areas at protecting birds by taking advantage of a large dataset of bird counts across a near-continental area – the North American Breeding Bird Survey (Pardieck et al., 2017). Controlling statistically for altitude and productivity in order to reduce the effect of PA location biases, we estimated PA effectiveness on two levels of biodiversity: on species’ assemblages, through indices of richness and summed abundance; and on individual common species, by estimating the effect of PAs on species’ abundance. At the assemblage level, we expected to find higher species diversity and abundance inside PAs. Indeed, as human activities are causing species population declines and local extinctions (Ceballos et al., 2017), and as PAs are expected to buffer against these activities, this should predictably lead to overall higher species richness and higher total abundance inside PAs, as found by Coetzee et al. (2014) and Gray et al. (2016). At the individual species level, we expected higher abundances within PAs. However, given differences in species’ habitat requirements, this result cannot be expected to hold universally (*i.e.,* species are not all expected to be more abundant in all PAs). For example, we expected protected forests to have a positive effect on forest species, but not on grassland species. Hence, we controlled in our analyses for broad vegetation structure (forest, shrub, herbaceous), by investigating separately the effects of PAs dominated by a particular vegetation structure on species with different habitat requirements. Additionally, we expected species with overall declining populations (thus more affected by anthropogenic activities), and species that avoid human presence (more sensitive to human disturbance) to present higher abundances inside PAs.

## Methods

In this study, we use the term “PA effectiveness” to refer to the difference in species diversity or abundance between protected and unprotected sites. This difference combines the effects of PA site selection (i.e., differences existing prior to the implementation of PA, for example if they are implemented in sites with higher-than-average richness or abundances) and the effects of protection itself (i.e., difference that arise after PA designation, if PAs effectively reduce population depletions and species local extinctions).

### Bird data

We used data from the North-American Breeding Bird Survey (BBS), a long-term volunteer-based monitoring scheme in Canada, the USA, and Mexico (Pardieck et al., 2017), version 2016.0. Our study region encompasses solely Canada and the USA, as few Mexican routes are monitored. This program is based on the annual monitoring of thousands of routes, each approximately 40 km long, during the bird breeding season. Each route is split into 50 stops; at each stop, the observer counts every bird heard or seen during three minutes, before moving to the next stop.

Given the length of BBS routes, they often intersect multiple land use types (e.g. forested, urban, agriculture; each with different bird assemblages), and they are rarely wholly contained within protected sites (most of the routes that cross PAs do so only in small fractions of their length). As a result, whole BBS routes are not particularly suited sampling units for investigating how PAs affect bird species. We chose instead to focus on small sections of routes – sequences of five stops, covering about 4 km – in order to obtain field-sampling units that are more homogeneous in terms of land use types and whose bird assemblages can be more directly linked to local landscape characteristics, especially protection. For each route, we only used the first sequence of five stops, because the only precisely georeferenced point we had access to was the first stop of each route. Indeed, even if in principle additional stops are spaced about 0.8 km from each other, in practice this distance can vary, making the location of additional stops in each route progressively more imprecise. Henceforth, and for simplicity, we use the term “routes” to refer to these initial sections of five stops rather than to entire BBS routes.

We excluded aquatic and nocturnal taxa, which are not well detected by this diurnal road-based monitoring scheme, as well as hybrid individuals. We also excluded seven non-native species, as they are not the focus of conservation efforts. The main dataset we analysed included 400 species in total. To test if removing non-native species can bias analyses (e.g, because they replace native species) we also ran analyses including these species (results are presented in Appendix S5).

Following Kendall et al. (1996)’s recommendations, we removed from the dataset the first year of participation of each BBS observer, to reduce bias due to differences in observer experience. To do so, we extracted the observers’ identifying number from the “Weather” file of the dataset (Pardieck et al., 2017) and calculated for each observer the first year of data collection reported in the dataset. We then removed every observation made by this observer this given year. We also removed double counts, which can be either due to the presence of two observers or an observer sampling several times in one year, by excluding observations for which the ‘RPID’ code (i.e. Run Protocol type) was 102, 103, 104, 203 or 502. We then focused on routes sampled at least 5 years between 2007 and 2016, obtaining a set of 3,046 routes. For routes sampled more than five years, we analysed only five (randomly selected) years of data, thus ensuring a consistent sampling effort across all routes. We fixed this arbitrary threshold of five years as a trade-off between obtaining high data quantity (number of routes analysed) and data quality (number of species detected per route, which increases with the number of years sampled). For each species, counts were summed across the five points and the five years, to obtain a single value per species per route, which we used as a measure of abundance. We acknowledge that these values correspond only to detected birds rather than true abundances. Detection is known to vary between habitats, depending on vegetation structure (Pacifici et al., 2008). This could lead to a difference of detection probability (and thus of perceived abundance) between protected and unprotected sites if vegetation structure differs; controlling for vegetation structure in our analyses should reduce this bias.

### Landscape data

For each route, we analysed the properties of the landscape within a 500 m buffer around the route’s 4 km track (total area ca. 5 km^2^). Given that 500 m corresponds broadly to the bird detection radius of the BBS (Sauer et al., 2017), we considered this a suitable description of the environment affecting the composition of birds detected by the BBS. Small et al. (2012) found that the immediate landscape composition (buffer of 0.4 km) of BBS routes was similar to the large-scale landscape composition (buffer of 10 km), so we do not expect this choice to strongly affect the results.

A Protected Area is defined by the IUCN as “a clearly defined geographical space, recognised, dedicated and managed, through legal or other effective means, to achieve the long term conservation of nature with associated ecosystem services and cultural values” (UNEP-WCMC IUCN, 2016). PAs are categorised by the IUCN within seven categories based on their protection level. At one extreme, Ia PAs are “strictly protected areas set aside to protect biodiversity […], where human visitation, use and impacts are strictly controlled and limited to ensure protection of the conservation values”. At the other extreme, VI PAs “conserve ecosystems and habitats together with associated cultural values and traditional natural resource management systems [and] are generally large, with most of the area in a natural condition, where a proportion is under sustainable natural resource management and where low-level non-industrial use of natural resources compatible with nature conservation is seen as one of the main aims of the area” (UNEP-WCMC and IUCN, 2018). We used data on locations (polygon shapefle) and IUCN categories of PAs collated in the World Database on Protected Areas (UNEP-WCMC and IUCN, 2018). According to the Word Database of Protected Areas methodology to calculate area covered by PAs (UNEP-WCMC and IUCN, 2019), we excluded “Man and Biosphere” reserves and PAs for which implementation was not finalised, keeping only PAs with a status “designated”, “inscribed” or “established”. In addition, we also removed PAs that were not spatialized (no polygon associated). Using QGis (QGIS Development Team, 2017), we calculated the proportion of area inside each route’s buffer that falls within a PA (all IUCN-categories combined, and dissolved to avoid double-counting of areas under multiple PA designations). We have also run analyses considering stricter PAs only (categories I-IV), as the effectiveness can vary with protection level (Gray et al., 2016; Coetzee et al., 2014).

We characterised each route according to four additional landscape variables, using QGis (QGIS Development Team, 2017): net primary productivity, altitude, human footprint, and type of vegetation structure. The first three are continuous variables, available as raster files, and we obtained a value per route by calculating the mean value across all pixels that overlap the respective buffer. We calculated net primary productivity as the mean during spring months (March to June) between 2004 and 2015 according to the monthly Net Primary Productivity Terra/Modis (NASA, 2017; resolution 0.1 degree, about 62 km^2^ at 45°N). Altitude was obtained from the GLOBE Digital Elevation Model (National Geophysical Data Center, 1999; resolution 0.008 degree, about 0.40 km^2^ at 45°N). Human footprint was derived from the 2009 Global terrestrial Human Footprint map (Venter et al., 2016; resolution 0.01× 0.008 degrees, about 0.50 km^2^ at 45°N). We defined the vegetation structure as a categorical variable with three types: forest, shrub and herbaceous. We used the Climate Change Initiative – Land Cover layer, using 2011 values as this is the central year of our sampling period (ESA, 2015, resolution 300m) and reclassified the land cover classes into the three vegetation structure types: forest from land cover classes 50-90 and 160-170 (N=1,282 routes; 97 protected by 50% at least); shrub, 120-122 (N=298, 30 protected); herbaceous, 130-153 (which includes croplands; N=1,214; 19 protected). We then obtained the main vegetation structure type for each route as the dominant one in the buffer. Routes dominated by other land use classes (mosaic, [30-40, 100-110 and 180]; bare areas [200-202]; water bodies [210]; urban [190], other [220]) were not analysed because they were too scarce. The 2,794 routes used in analyses are mapped in Appendix S1.

### Statistical analyses

We estimated the effect of PAs on each of two assemblage-level indices (species richness and summed abundance) and on the abundance of individual species using General Additive Models (GAMs). Models all had identical structures, with the response variable modelled as a function of a one-way interaction between the proportion of PAs inside the buffer and the type of vegetation structure. We added smoothed terms controlling for productivity and altitude, as well as longitude and latitude in order to correct for spatial autocorrelation. The general structure of the GAMs was: Response ~ PA * vegetation + s(productivity, altitude, longitude, latitude)

### Assemblage-level analyses

For each route, and across all 400 bird species analysed, we calculated two assemblage indices, in each case using the cumulative number of species or individuals seen across the 5 stops, over 5 years: species richness (*μ* = 28.5 ± 9.4 species); summed bird abundance across all species (*μ*= 248 ± 90 individuals). We then used a GAM to model each of these two assemblage variables against the above-mentioned covariates, assuming a Gaussian distribution for richness and a negative binomial distribution for abundance.

### Species-level analyses

We excluded the rarest species from this analysis in order to have enough statistical power, keeping only the 133 species observed in at least 150 routes. Under this threshold, numerous species were too rarely detected within protected routes, leading to aberrant estimates of PA effectiveness (either highly positive or highly negative). For each species, we only analysed routes that fall within the species’ distribution within our study area. We defined this distribution as the 90 % spatial kernel of the routes where the species was observed, obtained using the ‘*adehabitat*’ R package (Calenge, 2006). We treated all routes inside the kernel where the species was not observed as having zero abundance.

We modelled each species’ abundance using a GAM as mentioned above, with a negative binomial distribution. We then calculated for each species a “PA effect” (PAE), measured as the difference in predicted abundance between a fully protected (i.e. 100 % protected) and an unprotected route (i.e. 0 % protected) with all control variables fixed to their median values. We calculated PAE separately for each of the three types of vegetation structure, to obtain for each species a value of PAE_For_ for routes dominated by forest, PAE_Shrub_ for shrub routes, and PAE_Herb_ for herbaceous routes.

For each type of vegetation structure, we studied PAE values in order to understand the factors explaining which species are favoured or not by PAs. To do so, we used a linear model and a phylogenetic linear model with similar structures using species-level covariates. We considered three covariates: species’ habitat preference, population trend, and human-affinity. We used species’ main habitat compiled in 11 classes by Barnagaud et al. (2017, see Fig.2). We used species’ population trends in North America between 1966 and 2015, calculated for each species by Sauer et al. (2017) from the BBS data (negative numbers for declining species, positive for increasing species). We winsorized these values, folding down the 2.5% extreme values on each side, bringing estimates to a Gaussian distribution. Finally, we estimated for each species a human-affinity index, as the median human footprint of the routes where the species was observed, weighted by species’ abundance on the route. This index was calculated across all 3,046 routes prior to the exclusion of routes based on habitat types (i.e. also including routes dominated by mosaic, bare areas, water bodies and urban land cover) to be more representative of the diversity of habitats where species are present.

**Figure 1:**
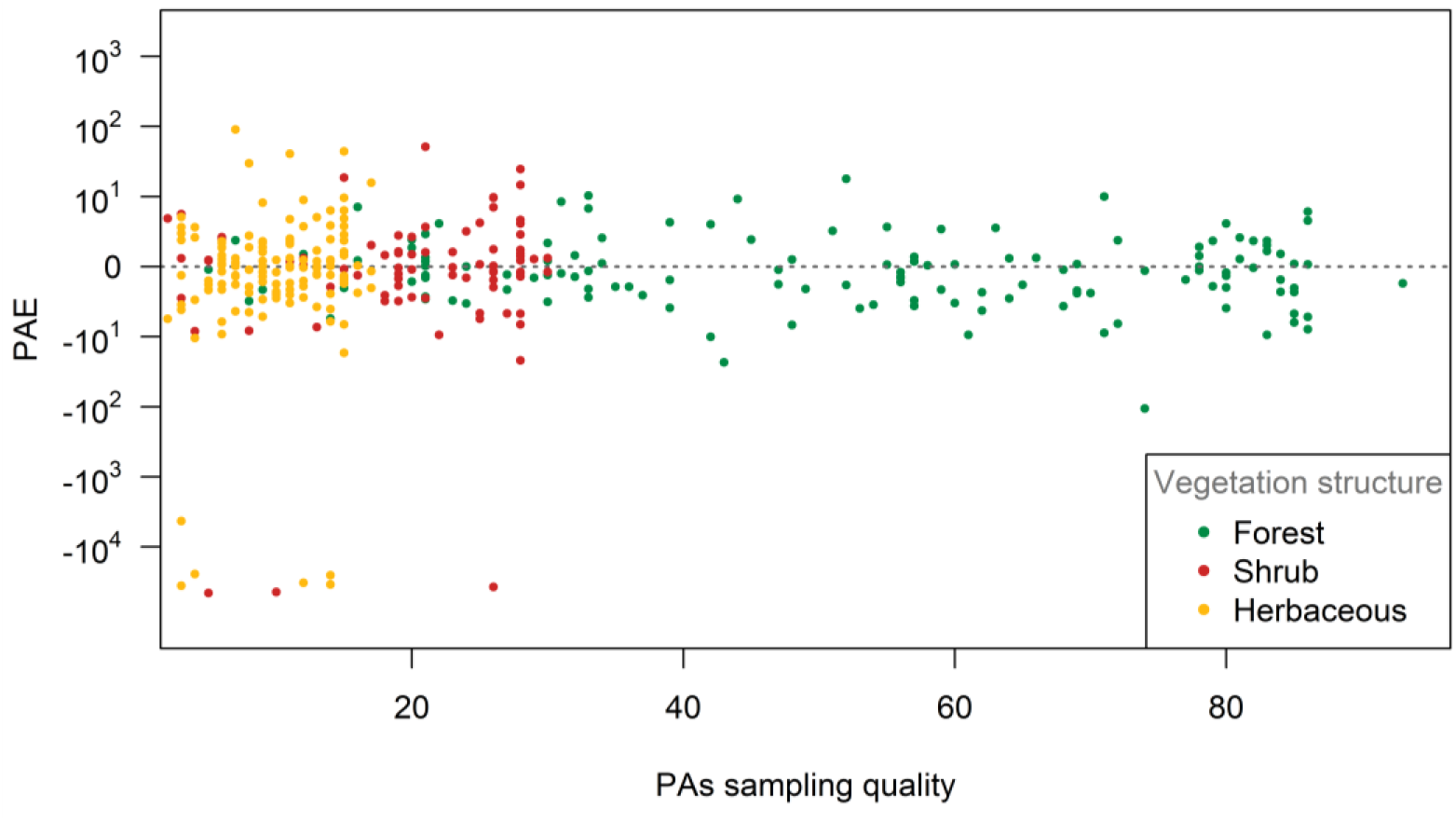
Estimated Protected Areas effect per species (PAE) (represented on a log scale in both negative and positive values), against PAs sampling quality, per vegetation structure type of the routes. PAs sampling quality is the number of routes within the species ‘ kernel with at least 50% of the buffer area covered by Protected Areas. Each point in the plot corresponds to a species, and each species can be represented by up to three points, one for each vegetation structure type.

**Figure 2:**
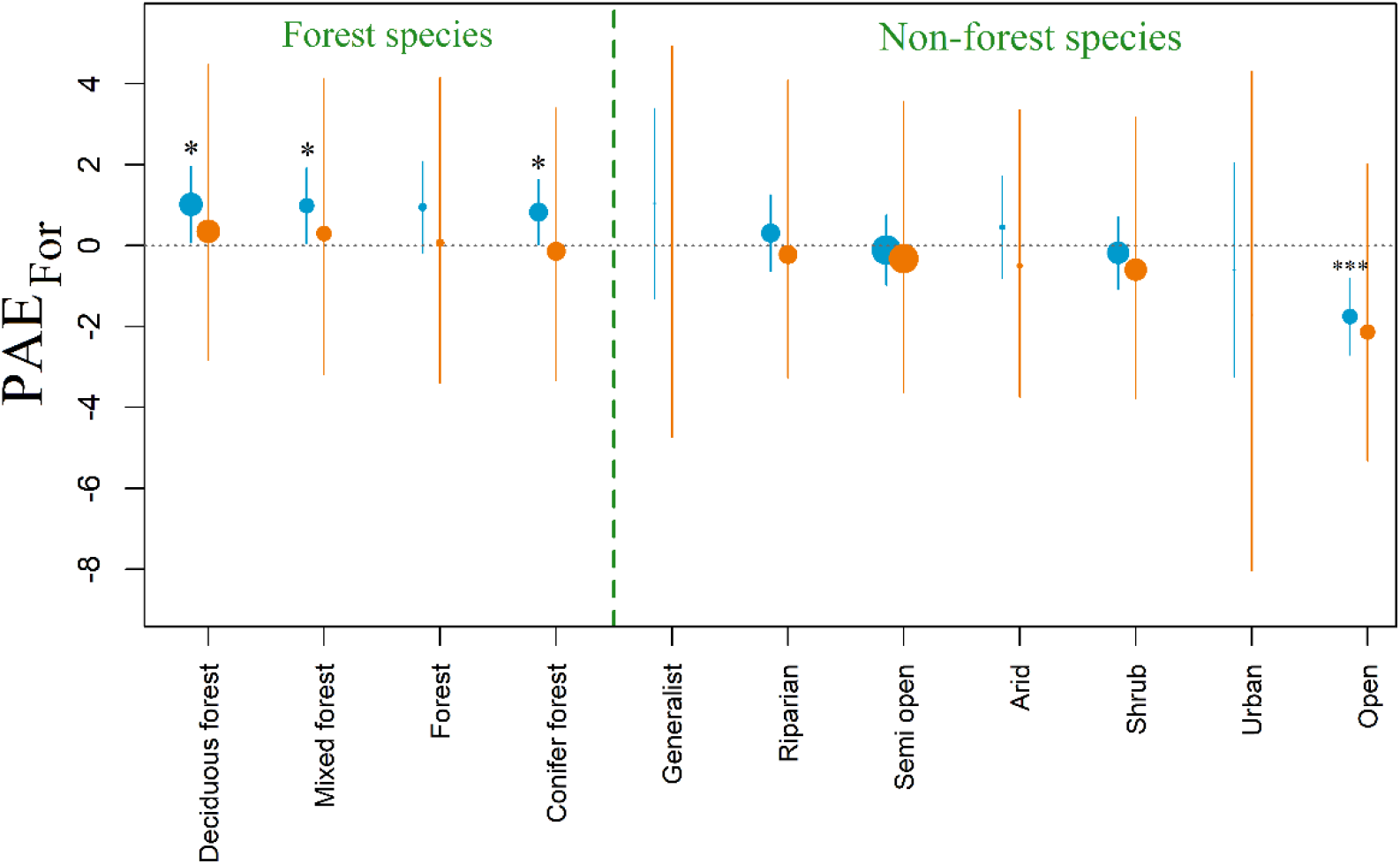
Estimated effect of Protected Areas on species within forest routes (PAE_For_) per species’ main habitat, estimated with a linear model (blue) and a phylogenetic linear model with Brownian motion (orange). Estimates were all calculated with species’ population trend and human-affinity fixed to zero. Error bars represent 95% CI; dot sizes are proportional to the number of species in each habitat group. Stars indicate significant effects for the particular model, for the particular species’ main habitat (P: 0.05 < * < 0.01 < ** < 0.001 < ***). Habitat types are ordered from the highest to the lowest PAE_For_ values under the phylogenetic model.

To account for phylogenetic autocorrelation, we ran a Brownian motion model implemented in the ‘*phylolm*’ R package (Tung Ho and Ané, 2014). To obtain the bird phylogeny, we selected randomly 100 phylogenetic Hackett backbone trees over 10,000 from Jetz et al. (2012) and calculated a maximum clade credibility tree using *Tree Annotator* from *Mr Bayes* (Drummond et al., 2012) with no burnin, and node heights calculated with the median. Confidence intervals of the phylogenetic model were estimated with the ‘*phylolm*’ function, using a bootstrap with 100 bootstrap replicates (Ives and Garland, 2010).

## Results

### Assemblage-level analyses

At the assemblage level, species richness and summed abundance differed significantly between vegetation structure types (respectively P=0.013, P=4.10^-10^), underlying the importance of accounting for habitat differences when studying PA effects.

Neither species richness nor summed abundance were significantly affected by the proportion of PAs in models that did not control for vegetation structure (respectively – 0.15 ± 0.72, P=0.835 and – 0.046 ± 0.031, P=0.143). In models controlling for vegetation structure, species richness did not vary significantly with protection within forest or within shrub routes (respectively – 1.39 ± 0.90, P=0.121 and – 0.305 ± 1.581, P=0.847) but increased with protection for herbaceous routes (4.35 ± 1.90, P=0.022). Summed abundance lightly decreased with protection within forest routes (−0.084 ± 0.039, P=0.030) but did not vary with protection within shrub or within herbaceous routes (respectively 0.082 ± 0.069, P=0.232 and 0.084 ± 0.082, P=0.307).

### Species-level analyses

According to the linear model, values of PAE_For_ – the predicted difference in a given species’ abundance between fully protected *versus* unprotected forest routes – differed significantly depending on the species’ main habitat. Hence, species that have forest as their main habitat showed higher abundances in protected than unprotected forests (Table 1, Fig.2). This difference was significant for the three main forest habitat preferences (i.e. mixed, deciduous, conifer) but not for the general forest category, which only represented five species (Table 1). Species favouring open habitats were significantly less abundant in protected forests than in unprotected forests (Table 1, Fig.2). We found no significant PA effect within forest routes for species favouring other habitat types. Species’ population trends between 1966 and 2015 did not significantly explain PAE_For_ (Table 1). In contrast, species’ human-affinity tended to be negatively correlated with PAE_For_ (*i.e.,* we found higher effects of PAs for species with lower affinity to humans in forested routes; Table 1, Fig.3). This trend was still present when we considered only forest species, but was not significant either (green dots in Fig.3; see Supporting Information in Appendix S2 for additional test).

**Table 1:** Model summaries regarding the estimated effect of Protected Areas on species within forest routes (PAE_For_): linear model and phylogenetic linear model with Brownian motion model. The top part gives estimates and P-values for all covariates, the bottom part gives estimates and P-values for all species’ habitat preferences, with trend and human-affinity fixed to zero. N corresponds to the number of species in each case.

|  | Linear model |  |  | Phylogenetic model |  |  |
| --- | --- | --- | --- | --- | --- | --- |
|  | *Estimate* | *SE* | *P* | *Estimate* | *SE* | *P* |
| Habitat | - | - | **4.10<sup>-6</sup>** | - | - | NA* |
| Trend | 2.10<sup>-3</sup> | 0.13 | 0.988 | 0.01 | 0.10 | 0.920 |
| Human-affinity | -0.08 | 0.05 | 0.087 | 0.001 | 0.04 | 0.979 |
| Mixed forest (N=10) | 0.99 | 0.49 | **0.042** | 0.30 | 1.82 | 0.869 |
| Forest (N=5) | 0.95 | 0.59 | 0.105 | 0.06 | 1.84 | 0.974 |
| Deciduous forest (N=18) | 1.02 | 0.49 | **0.037** | 0.35 | 1.80 | 0.846 |
| Conifer forest (N=11) | 0.83 | 0.42 | **0.048** | -0.14 | 1.77 | 0.937 |
| Semi open (N=19) | -0.11 | 0.45 | 0.805 | -0.33 | 1.82 | 0.856 |
| Riparian (N=11) | 0.31 | 0.48 | 0.517 | -0.22 | 1.81 | 0.903 |
| Generalist (N=1) | 1.04 | 1.20 | 0.386 | 0.13 | 2.36 | 0.956 |
| Shrub (N=15) | -0.18 | 0.47 | 0.700 | -0.60 | 1.79 | 0.737 |
| Arid (N=2) | 0.45 | 0.65 | 0.487 | -0.50 | 1.86 | 0.788 |
| Open (N=11) | -1.76 | 0.50 | **0.0004** | -2.14 | 1.81 | 0.238 |
| Urban (N=1) | -0.60 | 1.364 | 0.660 | -1.72 | 3.70 | 0.642 |

**Figure 3:**
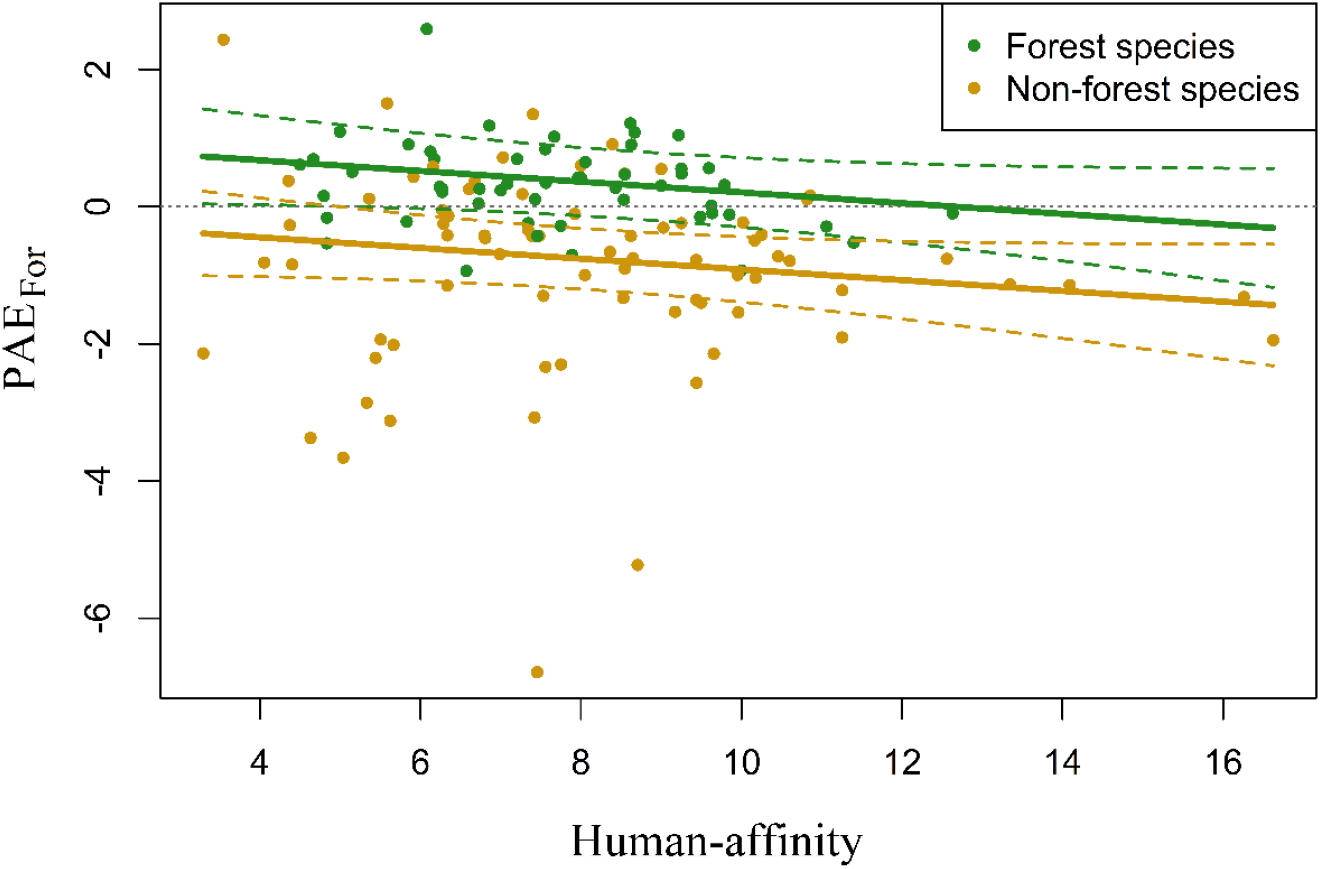
Effect of human-affinity (higher for species found preferably in areas of higher human footprint) on species’ responses to Protected Areas within forest routes (PAE_For_, above zero for species whose abundance in forest routes is higher in protected rather that in unprotected areas). Forest species (green) are species whose main habitat is “forest”, “conifer forest”, “mixed forest” or “deciduous forest”; non-forest species (brown) are all other species. Lines correspond to the effect of human-affinity on PAE_For_ for deciduous forest species (green) and semi-open species (brown), predicted by the linear model and their 95% confidence intervals in dashed lines.

None of these effect was significant in the phylogenetic model (Table 1 and Fig.2), which presented large confidence intervals.

The effect of PAs within shrub routes (PAE_Shrub_) was not affected by species’ main habitat and was only significantly negatively correlated with human-affinity (Supporting information, Table S1). PAE_Herb_ was only significantly affected by habitat preferences, being negative for conifer-forest species (Supporting information, Table S2). These results, however, need to be interpreted taking into account that shrub or herbaceous protected routes were rare in our dataset: on average, each species’ kernel included only 13 shrub and 9 herbaceous routes protected by 50% or more, in contrast with 50 protected forest routes (Fig.1; see Appendix S1 and S7 in Supporting Information).

Results, both at the assemblage and at the species levels, were similar but less significant when we considered only PAs of stricter management, as defined by IUCN categories I-IV (Dudley, 2008; see Supporting Information, Appendix S6). For shrub and herbaceous routes, the number of protected routes was even smaller than when all PAs were considered, leading to outlier results. Including non-native species in the analyses had little effect on the results at the assemblage level, the only difference being that the effect of protection within herbaceous routes became no longer significant (Appendix S5). At the species level, adding the three non-native species detected on more than 150 routes did not change the results (Appendix S5).

## Discussion

We compared the effect of PA coverage on bird species diversity, using assemblage indices (species richness, summed abundance) and individual species’ abundances.

At the assemblage level, we found very little effect of protection, only a small increase in richness in herbaceous routes and a small decrease in overall abundance in protected routes. In one sense, this is not surprising, as several large-scale studies found that assemblage metrics – particularly species richness – are relatively resilient to disturbance through species substitution (Dornelas et al., 2014; Supp and Ernest, 2014). Moreover, areas with low human-induced disturbance can have higher species richness than a pristine area, as predicted by the intermediate disturbance hypothesis (Roxburgh et al., 2004). Accordingly, Hiley et al., (2016) found lower alpha avian diversity in Mexican PAs than in unprotected areas. However, our results contrast with previous studies investigating this question such as Coetzee et al. (2014) or Gray et al. (2016), who found a positive effect of PAs on species richness and on summed abundance, including in North America (Coetzee et al., 2014). These two studies being meta-analyses, it is possible that a publication bias against studies showing negative or null effects of PAs (discussed by Coetzee et al., 2014) artificially increased the difference they measured. This is even more so given that the underlying studies of the meta-analyses were often designed to measure the effect of anthropogenic pressures, using PAs as benchmarks, rather than to measure the effectiveness of PAs (e.g. Sinclair et al., 2002; Bihn et al., 2008; Wunderle et al., 2006, all used in Coetzee meta-analysis), and may thus have focused on particularly intact protected sites and/or in highly degraded non-protected sites. Conversely, our results are not necessarily generalizable to other regions or other taxa, for example if North American birds are less sensitive to human activities than other taxa in North America or than birds in other regions, or if there is less contrast in human impacts in protected *versus* unprotected areas in North America than elsewhere. In our study, the observed lack of difference between protected and unprotected sites in terms of richness and abundance may also be an artefact of differences in species’ detectability (Boulinier et al., 1998), if PAs protect mainly species that are difficult to detect. This detection problem should not affect our result at the species level.

Even when overall species richness and abundance are similar, PAs may nonetheless have an effect on avian assemblages if different species respond differently to protection. We were only able to investigate this in depth for routes whose vegetation was dominated by forest, for which there was adequate sampling quality in PAs (Fig. 1). We found that among forest routes, PAs have an overall positive effect on species’ abundance, but only for those species with forest as their main habitat. In contrast, abundances of species favouring open habitats are negatively correlated with protection in forests. Forest PAs thus maintain a more forest-typical bird assemblage than comparable unprotected forests. These effects were significant with the linear model, but not the phylogenetic model. This suggests that much of the effect attributed to habitat preferences under the linear model relies on phylogenetic relatedness among species, which is not surprising as bird habitat preferences and phylogeny are correlated.

Contrary to our prediction, we did not find that species with low human-affinity (*i.e.,* species that avoid human-impacted areas) are significantly favoured by forest PAs, even if there was as nonsignificant positive effect. Also contrary to our expectation, and to previous results for common French birds (Devictor et al., 2007), we found no correlation between species’ population trends over the past 50 years and PAE_For_. This may reflect the fact that our model included only relatively common species (*i.e.,* observed on at least 100 routes in the studied years). It is thus possible that the most human-averse and endangered species are favoured by PAs, but that we could not measure it.

Our results suggest that PAs in herbaceous areas have a negative impact on conifer-forest species and on those with low human-affinity, whereas the effect of PAs in shrub routes was negatively correlated with human-affinity. Given the scarcity of protected routes within both of these vegetation structure types, we do not consider these results particularly robust or informative of the effectiveness of PAs, but they nonetheless emphasise the biases of BBS routes against shrub areas and herbaceous PAs (Appendix S7).

Given that PAs located in forests are not expected to favour the same species as PAs located in grasslands or shrub lands, we controlled for vegetation structure in our analyses of PA effects. However, this control masked the effect PAs may have had in preventing changes in vegetation structure (and associated changes in bird assemblages). For instance, given the vegetation structure categorisation we applied, the counterfactual for a protected forest was an unprotected forest, which does not take into account the possibility that the PA may have prevented the forest from being cleared. In other words, our approach does not measure the effect PAs can have on species diversity by preventing habitat destruction (that modifies vegetation structure type). Instead, it only measures the effects PAs can have in preventing habitat degradation (not modifying the vegetation structure type), for example from natural forest to exploited forest, or from natural grassland to croplands.

Pairwise comparisons of protected *versus* unprotected sites, and thus the meta analyses by Geldmann et al. (2013), Coetzee et al. (2014) and Gray et al. (2016), can integrate the combined effects of habitat destruction and habitat degradation on species diversity, given that the counterfactual chosen may have a different habitat structure from the protected site (e.g., a protected forest compared with an unprotected cropland). Nonetheless, because these metaanalyses build from underlying studies with a diversity of criteria in the choice of the counterfactuals, it is not straightforward to interpret the effectiveness values obtained. For instance, as discussed before, numerous underlying studies compared a highly degraded site with a protected site used as benchmark, with the purpose of estimating the impact of anthropogenic degradation, which can lead to an overestimate of PA effectiveness. Other studies aimed at estimating PA effectiveness directly (e.g. Wasiolka and Blaum, 2011; Lee et al., 2007), but their choice of counterfactual was subjectively based on what authors considered likely to have happened to the protected site had it not been protected (Coetzee et al., 2014). Finally, some other studies used in meta-analyses were not particularly interested in differences between protected and unprotected sites, with protection used only as a covariate to potentially explain variation around the signal the authors were interested in (e.g. Naidoo, 2004; McCarthy et al., 2010). So even though our approach does not allow us to measure the full effects of PAs, the differences we measured between protected and unprotected sites are defined statistically depending on the covariates included, which allowed to define more clearly how we measured PA effectiveness. A main advantage of using large biodiversity monitoring datasets (such as bird monitoring schemes) rather than pairwise comparisons is thus the possibility of applying a well-defined and repeatable control.

More broadly, our results emphasise that it is impossible to clearly measure the effectiveness of PAs in conserving species diversity without defining precisely what is expected from them. In this study, we measured PAs effectiveness as the difference in bird diversity between protected and unprotected sites, controlling for main landscape differences. This definition combines conservation ability to protect richest areas and to reduce effectively human pressures in these areas. If PAs are expected to present higher diversity in terms of assemblage metrics (species richness or summed abundance), then we found no evidence in our analyses that PAs are effective. If PAs are expected to protect all species’ populations, then we did not find they were effective either, as for about half of the 133 species studied here we found a negative effect of forest PAs. However, our results show that North-American forest PAs present higher abundances in forest species when compared with unprotected forest sites, and lower abundances of species favouring open habitats. That this result holds even though we found no significant difference in total abundance suggests that bird assemblages in protected forests are more forest-typical than those in unprotected forests. Our results thus indicate that forest PAs in North America are contributing to prevent forest habitat degradation, and associated losses in the abundance of forest specialist species. BBS routes do not currently cover sufficiently well other habitats besides forest to allow us to investigate whether the same result applies to PAs with a different vegetation structure, but datasets with a bigger proportion of sampling points inside PAs, across all habitats, would bring further light into this question.

## Data accessibility

All data used in the study (bird abundances, bird traits and landscape covariates) are publicly available, and can be obtained from the sources listed in the references.

## Supplementary material

Script and codes are available online: https://www.biorxiv.org/content/10.1101/433037v4.supplementary-material

## Supporting information

R Script and input files

## Acknowledgements

We thank Jean-Yves Barnagaud for his insightful comments and suggestions concerning the analyses. We are grateful to the thousands of U.S. and Canadian participants who annually perform and coordinate the Breeding Bird Survey.

This preprint has been peer-reviewed and recommended by Peer Community In Ecology (https://dx.doi.org/10.24072/pci.ecology.100018)

## Appendix

https://www.biorxiv.org/content/10.1101/433037v4.supplementary-material

