## Supplementary Information for "Using a large-scale biodiversity monitoring dataset to test the effectiveness of protected areas at conserving North-American breeding birds"

### SUPPORTING INFORMATION

#### Supplementary data

##### Appendix S1: Map of the routes used in the analyses

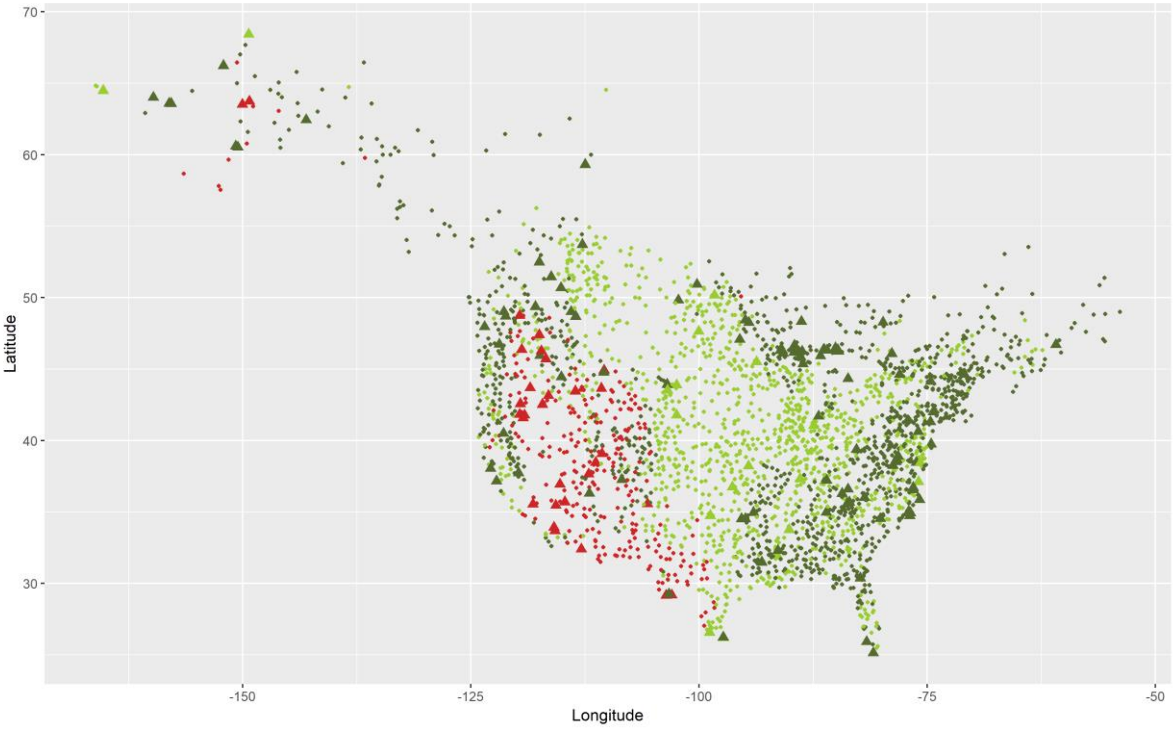

Fig.S1: Map of the 2,794 BBS routes used in the analyses. Colour of the points represents the main vegetation structure of the route (forest in dark green, herbaceous in light green and shrub in red). The shape of the point represents whether or not the route was protected by more than 50% (large triangle if protected by more than 50%, small point otherwise).

#### Supplementary results

##### Appendix S2: Human-affinity and $PAE_{For}$ for forest species only

As can be seen in Fig.3, the negative trend between the effect of Protected Areas on species within forest routes ($PAE_{For}$) and human-affinity is partly caused by a correlation between human-affinity and species habitat preferences (*i.e.*, forest species are more likely to have a low human-affinity than non-forest species). However, this trend still holds for forest species alone, suggesting that a forest species with low human-affinity is more likely to have a positive $PAE_{For}$ (*i.e.*, to be favoured by PAs) than a forest species with high human-affinity. This was checked by the following linear model with forest species only (N=53): $PAE_{For}$ ~ human-affinity. The result was nearly significant: −0.074 ± 0.045, P=0.10.

##### Appendix S3: Species model results for $PAE_{Shrub}$ and $PAE_{Herb}$

Table S1: Model summaries regarding the estimated effect of Protected Areas on species within shrub routes ($PAE_{Shrub}$) with the linear model and the phylogenetic linear model with Brownian motion model. The top part gives estimates and P-values for all covariates, the bottom part gives estimates and P-values for all species' habitat preferences, with trend and human-affinity fixed to zero. N corresponds to the number of species in each case. $PAE_{Shrub}$ values were winsorized to reduce the effect of extreme values, but some aberrant estimates remain, leading to high estimates in the model.

* P-values for the habitat variable as a whole could not be obtained, as Anova tables are not implemented in the *'phylolm'* package.

| | Linear model | | | Phylogenetic model | | |
|---|---|---|---|---|---|---|
| | *Estimate* | *SE* | *P* | *Estimate* | *SE* | *P* |
| Habitat | - | - | 0.447 | - | - | *NA** |
| Trend | 12 | 56 | 0.829 | -38 | 53 | 0.476 |
| Human-affinity | -52 | 23 | **0.027** | -67 | 21 | **0.0011** |
| Mixed forest (N=10) | 364 | 249 | 0.144 | 200 | 660 | 0.762 |
| Forest (N=5) | 336 | 370 | 0.365 | 295 | 703 | 0.675 |
| Deciduous forest (N=18) | -37 | 243 | 0.879 | 96 | 670 | 0.886 |
| Conifer forest (N=11) | 163 | 199 | 0.413 | 131 | 649 | 0.840 |
| Semi open (N=19) | 245 | 212 | 0.250 | 241 | 655 | 0.713 |
| Riparian (N=11) | 264 | 226 | 0.244 | 253 | 656 | 0.700 |
| Generalist (N=1) | 562 | 543 | 0.301 | 704 | 869 | 0.418 |
| Shrub (N=15) | 412 | 223 | 0.065 | 287 | 654 | 0.661 |
| Arid (N=2) | 383 | 296 | 0.196 | 371 | 677 | 0.584 |
| Open (N=11) | 381 | 234 | 0.104 | 253 | 664 | 0.703 |

Table S2: Model summaries regarding the estimated effect of Protected Areas on species within herbaceous routes ($PAE_{Herb}$) with the linear model and the phylogenetic linear model with Brownian motion model. The top part gives estimates and P-values for all covariates, the bottom part gives estimates and P-values for all species' habitat preferences, with trend and human-affinity fixed to zero. N corresponds to the number of species in each case. $PAE_{Herb}$ values were winsorized to remove the effect of extreme values.

* P-values for the habitat variable as a whole could not be obtained, as Anova tables are not implemented in the *'phylolm'* package.

| | Linear model | | | Phylogenetic model | | |
|---|---|---|---|---|---|---|
| | *Estimate* | *SE* | *P* | *Estimate* | *SE* | *P* |
| Habitat | - | | **0.011** | - | | *NA** |
| Trend | 0.19 | 0.24 | 0.426 | -0.18 | 0.30 | 0.547 |
| Human-affinity | 0.07 | 0.11 | 0.508 | 0.16 | 0.11 | 0.157 |
| Mixed forest (N=10) | -1.64 | 1.09 | 0.134 | -2.32 | 4.35 | 0.594 |
| Forest (N=5) | 0.20 | 1.31 | 0.879 | -1.80 | 4.42 | 0.684 |
| Deciduous forest (N=18) | -1.42 | 1.10 | 0.195 | -2.77 | 4.39 | 0.528 |
| Conifer forest (N=11) | -3.15 | 0.97 | **0.0011** | -4.38 | 4.33 | 0.312 |
| Semi open (N=19) | -0.16 | 1.03 | 0.877 | -0.67 | 4.37 | 0.878 |
| Riparian (N=11) | 0.03 | 1.33 | 0.982 | -0.19 | 4.46 | 0.966 |
| Generalist (N=1) | -2.37 | 2.77 | 0.392 | -2.86 | 5.74 | 0.618 |
| Shrub (N=15) | -0.37 | 1.04 | 0.722 | -0.27 | 4.39 | 0.951 |
| Arid (N=2) | -0.20 | 1.50 | 0.894 | 0.11 | 4.62 | 0.981 |
| Open (N=11) | -0.51 | 1.11 | 0.645 | -1.16 | 4.36 | 0.790 |
| Urban (N=1) | -5.85 | 3.14 | 0.062 | -8.18 | 8.98 | 0.362 |

**Appendix S4: List of 136 species used in the species level analysis. See methods for details about the column content.**

Non-native species included in the species analyses (see Appendix S5) are written in bold and are not included in the results presented in the main text.

| Scientific name | AOU | Species main habitat | Trend 1966-2015 | Human affinity | $PAE_{For}$ | $PAE_{Shrub}$ | $PAE_{Herb}$ |
|---|---|---|---|---|---|---|---|
| *Colinus virginianus* | 2890 | Open | -3.48 | 7.03 | 0.71 | 49.81 | 0.4 |
| *Phasianus colchicus* | 3091 | Open | -0.64 | 5.33 | -2.86 | 0.67 | 0.3 |
| *Meleagris gallopavo* | 3100 | Mixed Forest | 7.51 | 7.96 | 0.42 | -1.96 | 1.03 |
| *Zenaida macroura* | 3160 | Semi Open | -0.29 | 8.62 | -0.43 | -0.22 | -0.06 |
| *Buteo jamaicensis* | 3370 | Semi Open | NA | 7.55 | -0.6 | -0.07 | 0.66 |
| *Buteo lineatus* | 3390 | Deciduous Forest | 2.7 | 8.53 | 0.1 | -45461.88 | -1.03 |
| *Falco sparverius* | 3600 | Open | -1.39 | 8.71 | -5.22 | -0.24 | -2.37 |
| *Coccyzus americanus* | 3870 | Semi Open | -1.45 | 6.67 | 0.36 | 2.65 | 0.98 |
| *Picoides villosus* | 3930 | Forest | 0.81 | 5.83 | -0.22 | 1.73 | 2.34 |
| *Picoides pubescens* | 3940 | Deciduous Forest | 0.03 | 9.62 | 0.01 | -44010.13 | -1.91 |
| *Sphyrapicus varius* | 4020 | Deciduous Forest | 1.1 | 6.27 | 0.25 | 0.25 | 1.45 |
| *Dryocopus pileatus* | 4050 | Forest | 1.41 | 5.85 | 0.9 | NA | 2.49 |
| *Melanerpes carolinus* | 4090 | Forest | 1.02 | 9.48 | -0.15 | NA | -0.12 |
| *Colaptes auratus* | 4123 | Forest | NA | 6.51 | 0.13 | 1.02 | 0.74 |
| *Chaetura pelagica* | 4230 | Urban | -2.5 | 16.63 | -1.95 | NA | -5.17 |
| *Tyrannus forficatus* | 4430 | Semi Open | -0.78 | 8.39 | 0.91 | -2.83 | 0.46 |
| *Tyrannus tyrannus* | 4440 | Semi Open | -1.28 | 8.65 | -0.75 | -118.29 | -0.35 |
| *Tyrannus verticalis* | 4470 | Semi Open | 0.06 | 8.05 | -1 | -0.04 | 0.52 |
| *Myiarchus crinitus* | 4520 | Deciduous Forest | -0.03 | 9 | 0.3 | NA | -0.21 |
| *Myiarchus cinerascens* | 4540 | Shrub | 1.1 | 5.62 | -3.12 | 0.32 | -1.34 |
| *Sayornis phoebe* | 4560 | Deciduous Forest | 0.22 | 10 | -0.93 | -1904.58 | -1.73 |
| *Sayornis saya* | 4570 | Arid | 1.14 | 6.35 | -0.14 | 1.22 | 0.33 |
| *Contopus virens* | 4610 | Forest | -1.4 | 9.22 | 1.04 | NA | -0.45 |
| *Contopus sordidulus* | 4620 | Conifer Forest | -1.37 | 7.34 | -0.24 | 0.65 | -4.14 |
| *Empidonax virescens* | 4650 | Deciduous Forest | -0.26 | 8.62 | 1.21 | NA | -1.8 |
| *Empidonax traillii* | 4660 | Shrub | -1.48 | 11.25 | -1.91 | 0.96 | 0.95 |
| *Empidonax alnorum* | 4661 | Shrub | -0.89 | 7.48 | -0.43 | -0.96 | 0.16 |
| *Empidonax minimus* | 4670 | Semi Open | -1.71 | 5.36 | 0.11 | -8.25 | 2.39 |
| *Eremophila alpestris* | 4740 | Open | -2.46 | 5.44 | -2.2 | -1.98 | -2.44 |
| *Pica hudsonia* | 4750 | Semi Open | -0.49 | 7.42 | -3.07 | -0.65 | 0.47 |
| *Cyanocitta cristata* | 4770 | Mixed Forest | -0.66 | 11.06 | -0.29 | NA | -1.2 |
| *Corvus corax* | 4860 | Semi Open | 2.04 | 6.6 | 0.25 | -0.35 | -2.52 |
| *Corvus brachyrhynchos* | 4880 | Open | 0.07 | 10.18 | -1.04 | 0.92 | 0.06 |
| *Corvus ossifragus* | 4900 | Riparian | 0.48 | 16.26 | -1.32 | NA | 0.56 |
| ***Sturnus vulgaris*** | 4930 | Generalist | -1.46 | 14.85 | -2.85 | -0.60 | -1.01 |
| *Dolichonyx oryzivorus* | 4940 | Open | -2.06 | 7.45 | -6.79 | NA | -0.02 |
| *Molothrus ater* | 4950 | Semi Open | -0.66 | 7.39 | -0.43 | -0.63 | 0.86 |
| *Xanthocephalus xanthocephalus* | 4970 | Riparian | -0.06 | 3.54 | 2.43 | 0.39 | 1.85 |

| | | | | | | | |
|---|---|---|---|---|---|---|---|
| *Agelaius phoeniceus* | 4980 | Riparian | -0.93 | 9.44 | -1.36 | 0.53 | -0.16 |
| *Sturnella magna* | 5010 | Open | -3.28 | 9 | 0.54 | -26.26 | -0.27 |
| *Sturnella neglecta* | 5011 | Open | -1.29 | 4.63 | -3.37 | 0.35 | -1.38 |
| *Icterus spurius* | 5060 | Riparian | -0.87 | 8.55 | -0.91 | NA | 0.79 |
| *Icterus galbula* | 5070 | Semi Open | -1.49 | 9.65 | -2.15 | NA | -0.41 |
| *Icterus bullockii* | 5080 | Semi Open | -0.66 | 8.53 | -1.33 | 0.84 | 1.27 |
| *Euphagus cyanocephalus* | 5100 | Riparian | -2.25 | 7.53 | -1.3 | 0.76 | 0.11 |
| *Quiscalus quiscula* | 5110 | Semi Open | -1.75 | 12.56 | -0.76 | -1770.29 | -0.1 |
| *Haemorhous purpureus* | 5170 | Conifer Forest | -1.23 | 7.89 | -0.71 | 0.69 | -0.75 |
| *Haemorhous mexicanus* | 5190 | Arid | 0.12 | 13.35 | -1.14 | 0.72 | -1.32 |
| *Spinus tristis* | 5290 | Semi Open | -0.17 | 9.95 | -1 | -0.05 | 0.15 |
| *Pooecetes gramineus* | 5400 | Open | -0.85 | 5.04 | -3.66 | -0.38 | 0.6 |
| *Passerculus sandwichensis* | 5420 | Open | -1.36 | 7.75 | -2.3 | -0.1 | -0.2 |
| *Ammodramus savannarum* | 5460 | Open | -2.52 | 4.37 | -0.27 | 1.17 | 1.63 |
| *Chondestes grammacus* | 5520 | Semi Open | -0.78 | 5.66 | -2.02 | -1.17 | 0.56 |
| *Zonotrichia leucophrys* | 5540 | Semi Open | -0.4 | 6.33 | -1.15 | 1.28 | 0.75 |
| *Zonotrichia albicollis* | 5580 | Conifer Forest | -0.93 | 7 | 0.23 | 5.61 | -0.88 |
| *Spizella passerina* | 5600 | Semi Open | -0.6 | 10.82 | 0.11 | -0.02 | -0.21 |
| *Spizella pallida* | 5610 | Shrub | -1.14 | 5.5 | -1.94 | 1.24 | 1.89 |
| *Spizella breweri* | 5620 | Shrub | -1.01 | 3.29 | -2.14 | -0.27 | -0.93 |
| *Spizella pusilla* | 5630 | Semi Open | -2.33 | 9.44 | -2.57 | NA | 1.2 |
| *Junco hyemalis* | 5677 | Forest | NA | 6.67 | 0.49 | 0.15 | 0.57 |
| *Melospiza melodia* | 5810 | Shrub | -0.76 | 11.25 | -1.22 | 0.78 | 0.26 |
| *Melospiza lincolnii* | 5830 | Riparian | -0.36 | 5.91 | 0.43 | 1.32 | -0.99 |
| *Melospiza georgiana* | 5840 | Riparian | 0.93 | 8 | 0.6 | 0.6 | 2.76 |
| *Pipilo erythrophthalmus* | 5870 | Forest | -1.34 | 9.64 | -0.1 | NA | 0.73 |
| *Pipilo maculatus* | 5880 | Shrub | -0.03 | 7.34 | -0.34 | 0.22 | 2.99 |
| *Cardinalis cardinalis* | 5930 | Shrub | 0.32 | 10.6 | -0.79 | -3.96 | -0.45 |
| *Pheucticus ludovicianus* | 5950 | Forest | -0.86 | 7.08 | 0.33 | 0.33 | -0.53 |
| *Pheucticus melanocephalus* | 5960 | Deciduous Forest | 0.72 | 7.56 | 0.34 | 0.18 | -3.52 |
| *Passerina caerulea* | 5970 | Semi Open | 0.81 | 8.55 | 0.48 | 0.35 | 0.41 |
| *Passerina cyanea* | 5980 | Shrub | -0.73 | 9.03 | -0.31 | NA | 0.84 |
| *Passerina amoena* | 5990 | Arid | 0.21 | 4.4 | -0.84 | 0.36 | 2.98 |
| *Passerina ciris* | 6010 | Semi Open | -0.12 | 6.8 | -0.42 | -0.21 | 0.23 |
| *Spiza americana* | 6040 | Semi Open | -0.36 | 7.27 | 0.18 | 43.96 | 1.19 |
| *Piranga ludoviciana* | 6070 | Conifer Forest | 1.28 | 4.67 | 0.69 | 0.11 | 3.66 |
| *Piranga olivacea* | 6080 | Deciduous Forest | -0.22 | 8.62 | 0.9 | NA | 2.17 |
| *Piranga rubra* | 6100 | Mixed Forest | 0.22 | 7.2 | 0.69 | -50.97 | 0.9 |
| *Progne subis* | 6110 | Riparian | -0.91 | 10.46 | -0.73 | -2293.56 | -1.6 |
| *Petrochelidon pyrrhonota* | 6120 | Semi Open | 0.72 | 7.56 | -2.34 | -1.24 | -1.47 |
| *Hirundo rustica* | 6130 | Open | -1.19 | 9.96 | -1.54 | -0.61 | -0.02 |

| | | | | | | | |
|---|---|---|---|---|---|---|---|
| *Tachycineta bicolor* | 6140 | Riparian | -1.38 | 9.17 | -1.54 | -0.31 | 1.7 |
| *Tachycineta thalassina* | 6150 | Conifer Forest | -0.66 | 7.67 | 1.02 | -1.2 | 5.08 |
| *Stelgidopteryx serripennis* | 6170 | Riparian | -0.53 | 10.16 | -0.49 | -0.1 | -4.05 |
| *Bombycilla cedrorum* | 6190 | Mixed Forest | 0.07 | 12.63 | -0.1 | -7.29 | 0.35 |
| *Vireo olivaceus* | 6240 | Deciduous Forest | 0.75 | 8 | 0.42 | -2.41 | 0.97 |
| *Vireo gilvus* | 6270 | Deciduous Forest | 0.85 | 7.45 | -0.43 | -0.62 | -0.8 |
| *Vireo flavifrons* | 6280 | Deciduous Forest | 0.98 | 9.25 | 0.48 | NA | 0.34 |
| *Vireo solitarius* | 6290 | Mixed Forest | 2.86 | 6.17 | 0.69 | NA | -1.8 |
| *Vireo griseus* | 6310 | Riparian | 0.62 | 6.81 | -0.46 | -0.46 | 1.01 |
| *Mniotilta varia* | 6360 | Mixed Forest | -0.86 | 7.75 | -0.28 | NA | -2.75 |
| *Oreothlypis ruficapilla* | 6450 | Mixed Forest | 0.01 | 4.79 | 0.15 | -1.33 | 0.22 |
| *Oreothlypis celata* | 6460 | Shrub | -0.61 | 6.99 | -0.69 | 1.4 | 1.11 |
| *Setophaga americana* | 6480 | Conifer Forest | 1.11 | 7.43 | 0.11 | NA | -0.99 |
| *Setophaga petechia* | 6520 | Riparian | -0.61 | 8.36 | -0.66 | 1.76 | 1.18 |
| *Setophaga coronata* | 6556 | Conifer Forest | NA | 6.67 | 1.08 | 0.21 | -1.32 |
| *Setophaga magnolia* | 6570 | Conifer Forest | 0.87 | 6.57 | -0.94 | -0.94 | 0.16 |
| *Setophaga pensylvanica* | 6590 | Shrub | -1.15 | 7.92 | -0.11 | NA | 3.64 |
| *Setophaga fusca* | 6620 | Conifer Forest | 0.35 | 6.72 | 0.04 | 0.04 | -24359.41 |
| *Setophaga dominica* | 6630 | Conifer Forest | 0.98 | 8.55 | 0.47 | NA | -1.59 |
| *Setophaga virens* | 6670 | Mixed Forest | 0.35 | 6.24 | 0.29 | NA | 2.61 |
| *Setophaga pinus* | 6710 | Conifer Forest | 0.88 | 7.56 | 0.83 | NA | 1.31 |
| *Seiurus aurocapilla* | 6740 | Mixed Forest | -0.07 | 6.86 | 1.18 | 1.18 | -0.01 |
| *Parkesia noveboracensis* | 6750 | Riparian | 1.19 | 6.15 | 0.58 | -4.67 | 0.5 |
| *Geothlypis philadelphia* | 6790 | Shrub | -1.18 | 6.27 | -0.06 | -0.06 | -2.14 |
| *Geothlypis tolmiei* | 6800 | Conifer Forest | -1.66 | 4.83 | -0.54 | 1.49 | -4368.12 |
| *Geothlypis trichas* | 6810 | Riparian | -1.01 | 9.25 | -0.24 | 1.87 | 1.45 |
| *Icteria virens* | 6830 | Shrub | -0.62 | 6.29 | -0.25 | 2.01 | 2.29 |
| *Setophaga citrina* | 6840 | Deciduous Forest | 1.36 | 6.12 | 0.8 | NA | -10.44 |
| *Cardellina pusilla* | 6850 | Riparian | -1.8 | 6.33 | -0.42 | 1.54 | 3.85 |
| *Setophaga ruticilla* | 6870 | Deciduous Forest | -0.28 | 8.67 | 1.08 | -8.39 | -2.28 |
| ***Passer domesticus*** | 6882 | Generalist | -3.61 | 12.24 | -1.29 | -1.59 | -3.27 |
| *Mimus polyglottos* | 7030 | Semi Open | -0.46 | 10.02 | -0.23 | -0.76 | -0.11 |
| *Dumetella carolinensis* | 7040 | Shrub | -0.01 | 14.09 | -1.14 | -0.82 | 0.47 |
| *Toxostoma rufum* | 7050 | Shrub | -1.04 | 9.5 | -1.4 | NA | -0.27 |
| *Salpinctes obsoletus* | 7150 | Arid | -0.65 | 5.58 | 1.5 | 0.97 | -0.27 |
| *Thryothorus ludovicianus* | 7180 | Deciduous Forest | 1.04 | 9.85 | -0.12 | NA | -0.03 |
| *Thryomanes bewickii* | 7190 | Shrub | -0.9 | 7.4 | 1.34 | -0.98 | 2.3 |
| *Troglodytes aedon* | 7210 | Shrub | 0.26 | 9.43 | -0.78 | -0.36 | -0.16 |
| *Troglodytes hiemalis* | 7222 | Conifer Forest | 0.23 | 6.73 | 0.26 | 0.26 | -35800.29 |
| *Sitta carolinensis* | 7270 | Deciduous Forest | 1.71 | 9.25 | 0.55 | 2.62 | 1.02 |
| *Sitta canadensis* | 7280 | Conifer Forest | 0.72 | 5.15 | 0.5 | -0.14 | 2.11 |
| *Baeolophus bicolor* | 7310 | Deciduous Forest | NA | 9.93 | -0.39 | NA | -0.53 |

| | | | | | | | |
|---|---|---|---|---|---|---|---|
| *Poecile atricapillus* | 7350 | Deciduous Forest | 0.61 | 11.4 | -0.53 | 1.45 | 1.01 |
| *Poecile carolinensis* | 7360 | Mixed Forest | -0.38 | 9.79 | 0.31 | NA | -2.36 |
| *Poecile gambeli* | 7380 | Conifer Forest | -1.34 | 4.5 | 0.61 | 1.07 | -5.57 |
| *Regulus satrapa* | 7480 | Conifer Forest | -1.54 | 6.08 | 2.59 | -37373.47 | -32617.55 |
| *Regulus calendula* | 7490 | Conifer Forest | 0.47 | 4.83 | -0.17 | 0.84 | -25341.08 |
| *Polioptila caerulea* | 7510 | Deciduous Forest | 0.38 | 8.05 | 0.65 | 1.31 | 1.61 |
| *Hylocichla mustelina* | 7550 | Deciduous Forest | -1.91 | 9.6 | 0.56 | NA | 1.06 |
| *Catharus fuscescens* | 7560 | Deciduous Forest | -1.13 | 8.43 | 0.27 | -0.42 | -1.18 |
| *Catharus ustulatus* | 7580 | Mixed Forest | -0.84 | 6.27 | 0.21 | -1.37 | -0.81 |
| *Catharus guttatus* | 7590 | Mixed Forest | 0.33 | 4.99 | 1.09 | 0.05 | -34475.37 |
| *Turdus migratorius* | 7610 | Generalist | 0.12 | 10.86 | 0.16 | -0.24 | -1.58 |
| *Sialia sialis* | 7660 | Semi Open | 1.5 | 10.25 | -0.41 | -1928.64 | -0.81 |
| *Sialia currucoides* | 7680 | Semi Open | -0.54 | 4.05 | -0.82 | 0.32 | 2.4 |
| ***Streptopelia decaocto*** | 22860 | Semi Open | 29.18 | 9.68 | -1.06 | -0.58 | -5.17 |

#### Appendix S5: Assemblage analyses including non-native species

We ran the assemblage analyses without excluding non-native species to investigate if results were affected by the exclusion of these species. The 7 non-native species added to the assemblage analyses are: *Perdrix perdrix*, *Alectoris chukar*, *Columba livia*, *Sturnus vulgaris*, *Passer domesticus*, *Passer montanus*, *Streptopelia decaocto.*

<u>Assemblage models:</u>

Neither species richness, nor summed abundance were significantly affected by the proportion of PAs in the buffer in models not controlling for vegetation structure (respectively −0.62 ± 0.74, P=0.402 and −0.06 ± 0.032 P=0.064). In models controlling for vegetation structure, species richness did not vary significantly with protection in forest (−1.65 ± 0.91, P=0.07), shrub (−0.51 ± 1.61, P=0.752) or herbaceous routes (3.50 ± 1.95, P=0.073). Abundance did not vary significantly with protection shrub (0.068 ± 0.070, P=0.329) or herbaceous routes (0.04 ± 0.079, P=0.611) but decreased with protection in forest routes (−0.09 ± 0.04, P=0.026).

<u>Species models:</u>

Table S4: Model summaries regarding the estimated effect PAs on species within forest routes ($PAE_{For}$) with the linear model and the phylogenetic linear model with Brownian motion model. Three non-native species detected on at least 150 routes were included in this model (i.e. *Passer domesticus, Streptopelia decaocto, Sturnus vulgaris,* see Appendix S5). The top part gives estimates and P-values for all covariates, the bottom part gives estimates and P-values for all species' habitat preferences, with trend and human-affinity fixed to zero. N corresponds to the number of species in each case. This table is equivalent for Table 1, for strict PAs. For this model, we Winsorized PAEFor low values only (the 10% lowest values were pushed up to the value of the 10% quantile). This was necessary as the lack of power induced by the low number of protected routes with strict protection led to extreme estimates.

* P-values for the habitat variable as a whole could not be obtained, as Anova tables are not implemented in the *'phylolm'* package.

| | Linear Model | | | Phylogenetic Model | | |
|---|---|---|---|---|---|---|
| | *Estimate* | *SE* | *P* | *Estimate* | *SE* | *P* |
| Habitat | - | | **$2.10^{-6}$** | - | | *NA** |
| Trend | 0.015 | 0.10 | 0.882 | 0.023 | 0.12 | 0.849 |
| Human-affinity | -0.088 | 0.05 | 0.055 | -0.002 | 0.06 | 0.972 |
| Mixed forest (N=16) | 1.03 | 0.47 | **0.029** | 0.26 | 1.81 | 0.886 |
| Forest (N=7) | 0.99 | 0.57 | 0.083 | 0.05 | 1.73 | 0.977 |
| Deciduous forest (N=18) | 1.07 | 0.47 | **0.024** | 0.32 | 1.78 | 0.857 |
| Conifer forest (N=22) | 0.87 | 0.41 | **0.032** | -0.17 | 1.76 | 0.923 |
| Semi open (N=27) | -0.058 | 0.43 | 0.892 | -0.35 | 1.79 | 0.845 |
| Riparian (N=18) | 0.39 | 0.48 | 0.417 | -0.24 | 1.77 | 0.892 |
| Generalist (N=2) | -0.19 | 0.87 | 0.827 | -1.09 | 2.06 | 0.597 |
| Shrub (N=19) | -0.12 | 0.45 | 0.789 | -0.62 | 1.77 | 0.726 |
| Arid (N=5) | 0.49 | 0.63 | 0.439 | -0.53 | 1.83 | 0.772 |
| Open (N=14) | -1.44 | 0.51 | **0.0046** | -2.15 | 1.80 | 0.231 |
| Urban (N=1) | -2.62 | 1.42 | 0.065 | -1.64 | 3.67 | 0.655 |

#### Appendix S6: Equivalent analyses when considering PAs Ia-IV only

As the effectiveness of PAs may depend on the protection level they offer, we repeated did the exact same analyses as those presented in the main text, but considering only PAs categorised by the IUCN as categories Ia to IV (stricter conservation).

Assemblage models:

Neither species richness, nor summed abundance were significantly affected by the proportion of PAs in the buffer in models not controlling for vegetation structure (respectively −0.042 ± 0.092, P=0.665; −0.021 ± 0.050, P=0.692). In models controlling for vegetation structure, species richness did not vary significantly with protection in forest (−1.22 ± 1.19, P=0.305), shrub (−0.52 ± 1.92, P=0.787) but was increased by protection in herbaceous routes (8.04 ± 2.52, P=0.0014). Abundance did not vary significantly with protection in forest (−0.028 ± 0.052, P=0.590), shrub (−0.12 ± 0.08, P=0.153) or herbaceous routes (−0.16 ± 0.113, P=0.156).

Species models:

Table S4: Model summaries regarding the estimated effect of strict Protected Areas (IUCN categories Ia to IV only) on species within forest routes ($PAE_{For}$) with the linear model and the phylogenetic linear model with Brownian motion model. The top part gives estimates and P-values for all covariates, the bottom part gives estimates and P-values for all species' habitat preferences, with trend and human-affinity fixed to zero. N corresponds to the number of species in each case. This table is equivalent for Table 1, for strict PAs. For this model, we Winsorized PAEFor low values only (the 10% lowest values were pushed up to the value of the 10% quantile). This was necessary as the lack of power induced by the low number of protected routes with strict protection led to extreme estimates.

* P-values for the habitat variable as a whole could not be obtained, as Anova tables are not implemented in the *'phylolm'* package.

| | Linear Model | | | Phylogenetic Model | | |
|---|---|---|---|---|---|---|
| | *Estimate* | *SE* | *P* | *Estimate* | *SE* | *P* |
| Habitat | - | **-** | **$2.10^{-3}$** | - | - | *NA** |
| Trend | 0.009 | 0.11 | 0.932 | $4.10^{-4}$ | 0.11 | 0.997 |
| Human-affinity | -0.008 | 0.05 | 0.862 | 0.087 | 0.05 | 0.084 |
| Mixed forest (N=16) | 0.37 | 0.48 | 0.445 | -0.30 | 1.89 | 0.874 |
| Forest (N=7) | 0.34 | 0.60 | 0.571 | -1.03 | 1.94 | 0.596 |
| Deciduous forest (N=18) | 0.18 | 0.50 | 0.716 | -0.55 | 1.90 | 0.772 |
| Conifer forest (N=22) | 0.362 | 0.42 | 0.391 | -0.59 | 1.90 | 0.756 |
| Semi open (N=27) | -0.90 | 0.46 | **0.048** | -0.99 | 1.88 | 0.599 |
| Riparian (N=18) | -0.58 | 0.50 | 0.246 | -1.04 | 1.91 | 0.586 |
| Generalist (N=2) | 0.43 | 1.24 | 0.729 | -0.54 | 2.53 | 0.831 |
| Shrub (N=19) | -0.65 | 0.48 | 0.179 | -1.15 | 1.90 | 0.544 |
| Arid (N=5) | 0.40 | 0.68 | 0.554 | -0.50 | 1.95 | 0.798 |
| Open (N=14) | -1.44 | 0.51 | **0.0046** | -1.73 | 1.92 | 0.367 |
| Urban (N=1) | -2.62 | 1.42 | 0.065 | -4.16 | 3.92 | 0.289 |

#### Supplementary discussion:

##### Appendix S7: PA coverage differences between vegetation structure types in BBS routes

The better quality of sampling in forests results mainly from the fact that they are the most common vegetation structure type (49% of the studied area) and well covered by BBS routes (61% of the routes) and PAs (12% of the forests areas are protected and 7.6% of forest routes are protected by more than 50%). In contrast, shrub areas are the rarest vegetation structure type (14% of the studied area), they are even more rare in BBS routes (10% of the routes) but are well covered by PAs (15% of the shrub area are protected in the studied areas and 10.1% of the shrub BBS routes are protected by more than 50%). Herbaceous areas, which include cultivated areas, represent 34% of the studied area and 40% of BBS routes. Although they are relatively well covered by PAs in the studied area (10% protected), the first 5 stops of BBS routes do not intersect well with these herbaceous PAs as only 1.6% of the herbaceous routes are protected.
