## Supplementary material for "Using a large-scale biodiversity monitoring dataset to test the effectiveness of protected areas at conserving North-American breeding birds": R Script and input files: Change.Names.Species.R

```
vecteur<-replace(vecteur, vecteur=="Aimophila aestivalis", "Peucaea aestivalis")
vecteur<-replace(vecteur, vecteur=="Aimophila cassinii", "Peucaea cassinii")

vecteur<-replace(vecteur, vecteur=="Calcarius mccownii", "Rhynchophanes mccownii")

vecteur<-replace(vecteur, vecteur=="Stellula calliope", "Selasphorus calliope")
vecteur<-replace(vecteur, vecteur=="Carduelis flammea", "Acanthis flammea")
vecteur<-replace(vecteur, vecteur=="Carduelis lawrencei", "Spinus lawrencei")
vecteur<-replace(vecteur, vecteur=="Carduelis pinus", "Spinus pinus")
vecteur<-replace(vecteur, vecteur=="Carduelis psaltria", "Spinus psaltria")
vecteur<-replace(vecteur, vecteur=="Carduelis tristis", "Spinus tristis")

vecteur<-replace(vecteur, vecteur=="Carpodacus cassinii", "Haemorhous cassinii")
vecteur<-replace(vecteur, vecteur=="Carpodacus mexicanus", "Haemorhous mexicanus")
vecteur<-replace(vecteur, vecteur=="Carpodacus purpureus", "Haemorhous purpureus")

vecteur<-replace(vecteur, vecteur=="Dendragapus canadensis", "Falcipennis canadensis")

vecteur<-replace(vecteur, vecteur=="Dendroica caerulescens", "Setophaga caerulescens")
vecteur<-replace(vecteur, vecteur=="Dendroica castanea", "Setophaga castanea")
vecteur<-replace(vecteur, vecteur=="Dendroica cerulea", "Setophaga cerulea")
vecteur<-replace(vecteur, vecteur=="Dendroica coronata", "Setophaga coronata")
vecteur<-replace(vecteur, vecteur=="Dendroica discolor", "Setophaga discolor")
vecteur<-replace(vecteur, vecteur=="Dendroica dominica", "Setophaga dominica")
vecteur<-replace(vecteur, vecteur=="Dendroica fusca", "Setophaga fusca")
vecteur<-replace(vecteur, vecteur=="Dendroica graciae", "Setophaga graciae")
vecteur<-replace(vecteur, vecteur=="Dendroica magnolia", "Setophaga magnolia")
vecteur<-replace(vecteur, vecteur=="Dendroica nigrescens", "Setophaga nigrescens")
vecteur<-replace(vecteur, vecteur=="Dendroica occidentalis", "Setophaga occidentalis")
vecteur<-replace(vecteur, vecteur=="Dendroica palmarum", "Setophaga palmarum")
vecteur<-replace(vecteur, vecteur=="Dendroica pensylvanica", "Setophaga pensylvanica")
vecteur<-replace(vecteur, vecteur=="Dendroica petechia", "Setophaga petechia")
vecteur<-replace(vecteur, vecteur=="Dendroica pinus", "Setophaga pinus")
vecteur<-replace(vecteur, vecteur=="Dendroica striata", "Setophaga striata")
vecteur<-replace(vecteur, vecteur=="Dendroica tigrina", "Setophaga tigrina")
vecteur<-replace(vecteur, vecteur=="Dendroica townsendi", "Setophaga townsendi")
vecteur<-replace(vecteur, vecteur=="Dendroica virens", "Setophaga virens")
vecteur<-replace(vecteur, vecteur=="Parula americana", "Setophaga americana")

vecteur<-replace(vecteur, vecteur=="Oporornis formosus", "Geothlypis formosa")
vecteur<-replace(vecteur, vecteur=="Oporornis philadelphia", "Geothlypis philadelphia")
vecteur<-replace(vecteur, vecteur=="Oporornis tolmiei", "Geothlypis tolmiei")

vecteur<-replace(vecteur, vecteur=="Parus atricapillus", "Poecile atricapillus")
vecteur<-replace(vecteur, vecteur=="Parus carolinensis", "Poecile carolinensis")
vecteur<-replace(vecteur, vecteur=="Parus gambeli", "Poecile gambeli")
vecteur<-replace(vecteur, vecteur=="Parus hudsonicus", "Poecile hudsonicus")
vecteur<-replace(vecteur, vecteur=="Parus rufescens", "Poecile rufescens")

vecteur<-replace(vecteur, vecteur=="Pipilo crissalis", "Melozone crissalis")
vecteur<-replace(vecteur, vecteur=="Pipilo fuscus", "Melozone fusca")

vecteur<-replace(vecteur, vecteur=="Spizella arborea", "Spizelloides arborea")

vecteur<-replace(vecteur, vecteur=="Seiurus noveboracensis", "Parkesia noveboracensis")
vecteur<-replace(vecteur, vecteur=="Seiurus motacilla", "Parkesia motacilla")

vecteur<-replace(vecteur, vecteur=="Vermivora celata", "Oreothlypis celata")
vecteur<-replace(vecteur, vecteur=="Vermivora luciae", "Oreothlypis luciae")
vecteur<-replace(vecteur, vecteur=="Vermivora peregrina", "Oreothlypis peregrina")
vecteur<-replace(vecteur, vecteur=="Vermivora ruficapilla", "Oreothlypis ruficapilla")
vecteur<-replace(vecteur, vecteur=="Vermivora virginiae", "Oreothlypis virginiae")
vecteur<-replace(vecteur, vecteur=="Vermivora pinus", "Vermivora cyanoptera")

vecteur<-replace(vecteur, vecteur=="Wilsonia canadensis", "Cardellina canadensis")
vecteur<-replace(vecteur, vecteur=="Wilsonia citrina", "Setophaga citrina")
vecteur<-replace(vecteur, vecteur=="Wilsonia pusilla", "Cardellina pusilla")

vecteur<-replace(vecteur, vecteur=="Zoothera naevia", "Ixoreus naevius")
```
