## Supplementary material for "Using a large-scale biodiversity monitoring dataset to test the effectiveness of protected areas at conserving North-American breeding birds": R Script and input files: Complete script BBS.R

```
library(DescTools)
library(plyr)

'%not in%'<-function (x, table) is.na(match(x, table, nomatch=NA_integer_)) # Creates an exclusion function for subsets


setwd("D:/BBS")


###################
### Charge data ###
###################

# Charge the ten files of BBS data and merge the 10 tables into 1. Data are available here: https://www.pwrc.usgs.gov/bbs/RawData/
raw.data<-list()
for(i in 1:10) {raw.data[[i]]<-read.csv(file=paste0("RawData/fifty",i,".csv"), sep=",", header=T)}
BBS<-rbind(raw.data[[1]], raw.data[[2]] , raw.data[[3]] , raw.data[[4]] , raw.data[[5]], raw.data[[6]] , raw.data[[7]] , raw.data[[8]] , raw.data[[9]] , raw.data[[10]])
rm(raw.data)


### Data selection by year
an_min<-2007
an_max<-2016
nb_an<-an_max-an_min+1
bbs.raw<-subset(BBS, BBS$year>=an_min & BBS$year<=an_max)
bbs.raw$RtCode<-paste(bbs.raw$statenum, bbs.raw$Route, sep="_") # That is the unique identifying number I use in all analyses

### Charge "routes" file, available at: https://www.pwrc.usgs.gov/bbs/RawData/ 
routes.raw<-read.csv("routes.csv", sep=";", header=T)
routes.raw$RtCode<-as.factor(paste(routes.raw$statenum, routes.raw$Route, sep="_"))


### Remove first year of each observer, using weather table, available at: https://www.pwrc.usgs.gov/bbs/RawData/ 
weather<-read.csv("weather.csv")
weather$RteYear<-paste(weather$StateNum, weather$Route, weather$Year, sep="_")
bbs.raw$Observer<-weather[match(paste(bbs.raw$RtCode, bbs.raw$year, sep="_"), weather$RteYear), "ObsN"]
firstyear<-data.frame(Observer=levels(as.factor(bbs.raw$Observer)), First.Year=tapply(bbs.raw$year, as.factor(bbs.raw$Observer), min))
bbs.raw<-subset(bbs.raw, paste(bbs.raw$Observer, bbs.raw$year, sep=".") %not in% paste(firstyear$Observer, firstyear$First.Year, sep="."))

### Keep only 5 years per route to have consistent sampling effort between routes
# Remove routes monitored less than 5 years
rteparan<-subset(rteparan<-as.data.frame(table(bbs.raw$RtCode, bbs.raw$year)), rteparan$Freq>0)
frequents<-subset(frequents<-as.data.frame(table(droplevels(rteparan$Var1))) , frequents$Freq>=5)
rteparan<-subset(rteparan, rteparan$Var1 %in% frequents$Var1) 
rteparan$Var1<-droplevels(rteparan$Var1)
# Random selection of 5 years for routes monitored more than 5 years
for (i in 1:nlevels(rteparan$Var1)){
  sub<-subset(rteparan, rteparan$Var1==levels(rteparan$Var1)[i])
  if (nrow(sub)>5){aleat<-sample(paste(sub[,1], sub[,2]), (nrow(sub)-(nb_an/2))) # Identifies years that will be removed
  rteparan<-subset(rteparan, paste(rteparan$Var1, rteparan$Var2) %not in% aleat) 
  }}
Routes<-subset(routes.raw, routes.raw$RtCode %in% rteparan$Var1 & routes.raw$RouteTypeDetailId<4)    # Remove the routes too old, applies the removal following the random selection of 5 years 
bbs<-subset(bbs.raw, paste(bbs.raw$RtCode, bbs.raw$year) %in% paste(rteparan$Var1, rteparan$Var2) & bbs.raw$RtCode %in% Routes$RtCode) # Remove data associated with removed routes
table(table(bbs$RtCode, bbs$year)) # Check only fives


### Species selection
## Charge species table: available in appendix (we added a column for species removed from the analyses compared with the BBS-given version)
species<-read.csv("SpeciesList.VCa.csv", sep=";", header=T)
# Remove unwanted species
bbs<-subset(bbs, bbs$AOU %in% species$AOU[species$Removed %not in% c("aquatic", "hybrid", "unidentified", "Nocturnal")]) 
# Remove double counts (102-104, 502), double observers (203)
bbs<-subset(bbs, bbs$RPID %not in% c(102, 103, 104, 203, 502))
# Rollup subspecies to species using AOU
bbs$AOU<-replace(bbs$AOU, bbs$AOU %in% c(3380), 3370) # Buteo jamaicensis
bbs$AOU<-replace(bbs$AOU, bbs$AOU %in% c(4882, 4881), 4880) # Corvus brachyrhynchos
bbs$AOU<-replace(bbs$AOU, bbs$AOU %in% c(5660, 5690, 5670, 5680, 5671), 5677) # Junco hyemalis
bbs$AOU<-replace(bbs$AOU, bbs$AOU %in% c(6560, 6550), 6556) # Setophaga coronata
bbs$AOU<-replace(bbs$AOU, bbs$AOU %in% c(4120, 4125, 4130), 4123) # Colaptes auratus
# Add names
bbs$Name<-species[match(bbs$AOU, species$AOU), "Name"]
bbs$FName<-species[match(bbs$AOU, species$AOU), "Fr_Name"]


########## Create birds per route matrix
bbs$tot<-rowSums(bbs[,8:12]) # Sums abundances of the first five points of each route
bbs<-bbs[,-c(8:57)] # Removes other counts

obs<-data.frame(round(tapply(bbs$tot, list(bbs$RtCode, bbs$AOU), sum))) # Sum counts over years


################
### GIS WORK ###
################
library(sp) ; library(sf) ; library(shapefiles) ; library(raster) ; library(rgdal) ; library(rgeos)

# Charge buffers to use (calculated under QGis as the first 5 kilometers of BBS polylines available on the BBS website). 
buff.sf<-st_transform(read_sf("Buffer.to.use.SCR106.2019.shp"), st_crs("+proj=moll +lon_0=0 +x_0=0 +y_0=0 +ellps=WGS84 +units=m +no_defs"))


### Protected areas, shapefile available here: www.protectedplanet.net
shp.PA<-st_transform(read_sf("WDPA_Nov2018-shapefile-polygons.shp"), st_crs("+proj=moll +lon_0=0 +x_0=0 +y_0=0 +ellps=WGS84 +units=m +no_defs"))
shp.PA<-subset(shp.PA, shp.PA$STATUS %in% c("Designated", "Inscribed", "Established"))
shp.PA<-subset(shp.PA, shp.PA$DESIG_ENG != "UNESCO-MAB Biosphere Reserve")


# Intersect routes and PAs (considering either all PAs for main text, I-IV PAs for appendices)
PAs.interest<-st_intersection(buff.sf, st_buffer(shp.PA,0))
PAs.inter.TOT<-st_intersection(buff.sf,st_buffer(st_combine(PAs.interest),0))
PAs.inter.I.IV<-st_intersection(buff.sf,st_buffer(st_combine(PAs.interest[PAs.interest$IUCN_CAT %in% c("Ia", "Ib", "II", "III", "IV"),]),0))


## Add areas of routes (to divide the area occupied by PAs by the total area of buffers)
# Area of buffers
buff.sf$area<-st_area(buff.sf)
Routes$area.buff<-as.numeric(buff.sf$area[match(Routes$RtCode, buff.sf$RtCode)])

# For PAs tot
PAs.inter.TOT$area<-st_area(PAs.inter.TOT)
Routes$area.PA.TOT<-as.numeric(PAs.inter.TOT$area[match(Routes$RtCode, PAs.inter.TOT$RtCode)])
Routes$area.PA.TOT<-replace(Routes$area.PA.TOT, is.na(Routes$area.PA.TOT)==TRUE, 0)
Routes$prop.PA.TOT<-Routes$area.PA.TOT / Routes$area.buff
hist(Routes$prop.PA.TOT)

# For PAs I-IV
PAs.inter.I.IV$area<-st_area(PAs.inter.I.IV)
Routes$area.PA.I.IV<-as.numeric(PAs.inter.I.IV$area[match(Routes$RtCode, PAs.inter.I.IV$RtCode)])
Routes$area.PA.I.IV<-replace(Routes$area.PA.I.IV, is.na(Routes$area.PA.I.IV)==TRUE, 0)
Routes$prop.PA.I.IV<-Routes$area.PA.I.IV / Routes$area.buff
hist(Routes$prop.PA.I.IV)
plot(Routes$prop.PA.TOT, Routes$prop.PA.I.IV)


###### RASTERS
buff.ras<-spTransform(readOGR("Buffer.to.use.SCR43.shp"), CRSobj = CRS("+init=epsg:4238")) # Same shapefile than before, but different CRS and in sp format
buff.ras<-buff.ras[buff.ras$RtCode %in% Routes$RtCode, ]

### Altitude. Data available here: https://www.ngdc.noaa.gov/mgg/topo/gltiles.html
alt<-raster("globe_v1_all/w001001.adf")
alt<-crop(alt, extent(buff.ras), snap="out") # reduce range
buff.ras$alt<-extract(alt, buff.ras, fun=mean, df=T, na.rm=T)$w001001
Routes$Alt_mean<-as.numeric(buff.ras$alt[match(Routes$RtCode, buff.ras$RtCode)])

### NPP. Data available here: https://neo.sci.gsfc.nasa.gov/view.php?datasetId=MOD17A2_M_PSN
npp<-raster("NPP_seasonal_average_MarchJune_2004_2015.tif")
npp<-crop(npp, extent(buff.ras), snap="out") # reduce range
buff.ras$npp<-extract(npp, buff.ras, fun=mean, df=T, na.rm=T)$NPP_seasonal_average_MarchJune_2004_2015
Routes$NPP_mean<-as.numeric(buff.ras$npp[match(Routes$RtCode, buff.ras$RtCode)])

### HFP. Data available here: https://datadryad.org/resource/doi:10.5061/dryad.052q5
hfp<-raster("D:/Victor/PA GIS/Human Footprint/Footprint2.tif")
hfp<-crop(hfp, extent(buff.ras), snap=  "out")  # reduce range
buff.ras$hfp<-extract(hfp, buff.ras, fun=mean, df=T, na.rm=T)$Footprint2
Routes$hft<-as.numeric(buff.ras$hfp[match(Routes$RtCode, buff.ras$RtCode)])

### Land cover. Data available here: http://maps.elie.ucl.ac.be/CCI/viewer/index.php
glc<-raster("ESACCI-LC-L4-LCCS-Map-300m-P1Y-2011-v2.0.7.tif")
glc<-crop(glc, extent(buff.ras), snap="out")

# Group raster as indicated as: 991=Other, 992= For, 993=Shrub, 994=Herb, Mosaic=995, Bare=996 Water=997, Urban=998
transition<-matrix(c(c(0, 10, 11, 12, 20, 30, 40, 50, 60, 61, 62, 70, 71, 72, 80, 81, 82, 90, 100, 110, 120, 121, 122, 130, 140, 150, 152, 153, 160, 170, 180, 190, 200, 201, 202, 210, 220),
                     c(991,994,994,994,994,995,995,992, 992, 992,992,992,992, 992,992,992,992,992,995, 995,993,993,993,994,994,994,994,994,992,992,995,998,996,996,996,997,991) 
), ncol=2, nrow=37, byrow=F)

glc2<-reclassify(glc, transition)

# Extract the main land use for each buffer
buff.ras$main.glc<-extract(glc2, buff.ras, fun=function(x, na.rm=T){as.numeric(names(which(table(x)==max(table(x), na.rm=T))[1]))}, df=T, na.rm=T)$layer
Routes$mainoccG<-as.numeric(buff.ras$main.glc[match(Routes$RtCode, buff.ras$RtCode)])
Routes$mainoccG<-revalue(as.character(Routes$mainoccG), c("911"="Other", "992"="For", "993"="Shrub", "994"="Herb", "995"="Mosaic", "996"="Bare", "997"="Water", "998"="Urban"))


#############################################
### CALCULATE BIRD RICHNESS AND ABUNDANCE ###
#############################################
spcTot<-nlevels(as.factor(bbs$AOU)) # Total number of species in the data
B.index<-data.frame(RtCode=rownames(obs))


### WITH NON-NATIVE SPECIES
# Species richness
B.index$sp_richness.NONNAT<-apply(obs[,1:spcTot], 1, function(x) length(which(x>0))) 
Routes$sp_richness.NONNAT<-B.index[match(Routes$RtCode, B.index$RtCode), "sp_richness.NONNAT"]
hist(Routes$sp_richness.NONNAT)

# Abundance
B.index$abundance.NONNAT<-rowSums(obs[,1:spcTot], na.rm=T)
Routes$abundance.NONNAT<-B.index[match(Routes$RtCode, B.index$RtCode), "abundance.NONNAT"]
hist(Routes$abundance.NONNAT)


### WITHOUT NON-NATIVE SPECIES
# Remove species
write.csv(obs, "Obs.NNNative.BBS.Round1.11.01.19.csv") # Save obs file without removing introduced species
obs$X2881<-obs$X2882<-obs$X3131<-obs$X4930<-obs$X6882<-obs$X6883<-obs$X22860<-NULL # Removes the seven non-native species

# Species richness
B.index$sp_richness<-apply(obs[,1:(spcTot-7)], 1, function(x) length(which(x>0))) 
Routes$sp_richness<-B.index[match(Routes$RtCode, B.index$RtCode), "sp_richness"]
hist(Routes$sp_richness)

# Abundance
B.index$abundance<-rowSums(obs[,1:(spcTot-7)], na.rm=T)
Routes$abundance<-B.index[match(Routes$RtCode, B.index$RtCode), "abundance"]
hist(Routes$abundance)
plot(Routes$sp_richness, Routes$sp_richness.NONNAT) ; abline(a=0, b=1, col="red", lwd=2)
plot(Routes$abundance, Routes$abundance.NONNAT) ; abline(a=0, b=1, col="red", lwd=2)

# Extract obs without introduced species and routes
write.csv(obs, "Obs.BBS.Round1.11.01.19.csv")
write.csv(Routes, "Routes.BBS.Round1.11.01.19.csv", row.names=F)


###########################
### ASSEMBLAGE ANALYSES ###
###########################
library(mgcv)
library(visreg)
library(plyr)

obs<-read.csv("Obs.BBS.Round1.11.01.19.csv", row.names="X") ; obs<-obs[order(rownames(obs)),]
Routes<-read.csv("Routes.BBS.Round1.11.01.19.csv") ; Routes<-Routes[order(Routes$RtCode),]
table(rownames(obs)==Routes$RtCode) # Check 100% TRUE

Routes<-subset(Routes, Routes$mainoccG %in% c("For", "Herb", "Shrub"))
Routes$mainoccG<-factor(droplevels(Routes$mainoccG) , c("For","Herb","Shrub"))


### Main text
mA1<-gam(sp_richness ~ prop.PA.TOT * mainoccG + s(Alt_mean,NPP_mean,Latitude,Longitude), data=Routes, family=gaussian)
mA2<-gam(sp_richness ~ prop.PA.TOT            + s(Alt_mean,NPP_mean,Latitude,Longitude), data=Routes, family=gaussian)
mA3<-gam(abundance   ~ prop.PA.TOT * mainoccG + s(Alt_mean,NPP_mean,Latitude,Longitude), data=Routes, family=nb      )
mA4<-gam(abundance   ~ prop.PA.TOT            + s(Alt_mean,NPP_mean,Latitude,Longitude), data=Routes, family=nb      )


### Appendix I-IV
mB1<-gam(sp_richness ~ prop.PA.I.IV * mainoccG + s(Alt_mean,NPP_mean,Latitude,Longitude), data=Routes, family=gaussian)
mB2<-gam(sp_richness ~ prop.PA.I.IV            + s(Alt_mean,NPP_mean,Latitude,Longitude), data=Routes, family=gaussian)
mB3<-gam(abundance   ~ prop.PA.I.IV * mainoccG + s(Alt_mean,NPP_mean,Latitude,Longitude), data=Routes, family=nb      )
mB4<-gam(abundance   ~ prop.PA.I.IV            + s(Alt_mean,NPP_mean,Latitude,Longitude), data=Routes, family=nb      )


### Appendix non native species
mC1<-gam(sp_richness.NONNAT ~ prop.PA.TOT * mainoccG + s(Alt_mean,NPP_mean,Latitude,Longitude), data=Routes, family=gaussian)
mC2<-gam(sp_richness.NONNAT ~ prop.PA.TOT            + s(Alt_mean,NPP_mean,Latitude,Longitude), data=Routes, family=gaussian)
mC3<-gam(abundance.NONNAT   ~ prop.PA.TOT * mainoccG + s(Alt_mean,NPP_mean,Latitude,Longitude), data=Routes, family=nb      )
mC4<-gam(abundance.NONNAT   ~ prop.PA.TOT            + s(Alt_mean,NPP_mean,Latitude,Longitude), data=Routes, family=nb      )


##################################
### RUN SPECIES-LEVEL ANALYSES ###
##################################
library(adehabitat)
library(SDMTools)
library(MASS)
library(pscl)
library(mgcv)
library(DescTools)
library(plyr)


### SET THE CONDITIONS TO TEST
prot="I.IV" # Possibilities prot="TOT" (all protected areas) or "I.IV"  (strict protected areas)
espe="Nat"  # Possibilities espe="Nat" (only native species) or "NNNat" (both native and non-native species)


# Charge bird data file
if(espe=="Nat"){obs.full<-read.csv("Obs.BBS.Round1.11.01.19.csv", row.names="X") ; obs.full<-obs.full[order(rownames(obs.full)),]}
if(espe=="NNNat"){obs.full<-read.csv("Obs.NNNative.BBS.Round1.11.01.19.csv", row.names="X") ; obs.full<-obs.full[order(rownames(obs.full)),]}

Routes.full<-read.csv("Routes.BBS.Round1.11.01.19.csv") ; Routes.full<-Routes.full[order(Routes.full$RtCode),]
table(rownames(obs.full)==Routes.full$RtCode) # Check 100% TRUE

Routes<-subset(Routes.full, Routes.full$mainoccG %in% c("For", "Herb", "Shrub"))
Routes$mainoccG<-droplevels(Routes$mainoccG) 
obs<-obs.full[rownames(obs.full) %in% Routes$RtCode,] # Removes observation done on routes of other landuse than For, Herb, Shrub
table(rownames(obs)==Routes$RtCode) # Check 100% TRUE


# Remove rare species (they won't be modelled)
obs<-obs[,colSums(obs, na.rm=T)>0] # Remove routes with 0 obs (avoid buggs)
v<-NA ; for(i in 1:ncol(obs)){v[i]<-table(obs[,i]==0)[1]} # Calculate the number of routes of detection for each species
obst<-obs[,v >= 150] # Remove species detected in less than 150 routes

# Change NA for 0
obst[,grepl("X", names(obst))]<-replace(obst[,grepl("X", names(obst))], is.na(obst[,grepl("X", names(obst))])==TRUE, 0)

# Add coordinates in obs
obst$Lat<- Routes[match(rownames(obst), Routes$RtCode), "Latitude"]
obst$Lon<- Routes[match(rownames(obst), Routes$RtCode), "Longitude"]


# Apply Kernels
obsRed<-obst
sizeKer<-data.frame(AOU=as.numeric(rep("NA",ncol(obsRed))), size=as.numeric(rep("NA",ncol(obsRed))))

for (spc in 1:(ncol(obsRed)-2)){
  df<-data.frame(x=obst$Lon[obst[,spc]>0], y=obst$Lat[obst[,spc]>0]) # Create df with coordinates of routes where the species is present
  ver<-getverticeshr(kernelUD(df, id=rep("a", nrow(df))), 90) # Calculate Kernel, extract polygon of 90% Kernel
  Routes$Inside<-pnt.in.poly(cbind(Routes$Longitude, Routes$Latitude), cbind(ver$a[,2], ver$a[,3]))[,3] # Report whether each route is inside the Kernel polygon (1) or not (0)
  # Apply change in obsRed so points outside Kernel are NA for the species
  sub<-data.frame(RtCode=rownames(obst), OBS=obst[,spc])
  sub$INS<-Routes[match(sub$RtCode, Routes$RtCode), "Inside"] 
  vec<-sub$OBS/sub$INS # If the point is in the polygon, value is not changed (divided by 1), otherwise it is changed to NA (divided by 0) 
  vec<-replace(vec, vec=="Inf", NA) # Points outside the kernell but with the species -> Inf are removed (NA)
  obsRed[,spc]<-replace(vec, vec=="NaN", NA) 
  sizeKer$AOU[spc] <- substr((names(obsRed)[spc]), 2,10)
  sizeKer[spc,2] <- kernel.area(df, id=rep("a", nrow(df)))[15,]
}


# Add species habitats preferences
effects<-data.frame(AOU=substr(colnames(obsRed)[1:(ncol(obsRed)-2)],2,10))
traitsJYB<-read.table("LHT_USBirds4_Traits_JYB.txt", header=T) # Data available in Barnagaud et al. 2017
effects$J.main.habitat<-traitsJYB[match(effects$AOU, traitsJYB$AOU), "main_habitat"]
table(effects$AOU==substr(colnames(obsRed)[1:(ncol(obsRed)-2)],2,10)) # Check species in effects fit with names in obsRed (should be 100% TRUE)

obsRed<-obsRed[order(rownames(obsRed)),]
Routes<-Routes[order(Routes$RtCode),]
table(rownames(obsRed)==Routes$RtCode) # Check 100% TRUE

if(prot=="TOT"){Routes$PA<-Routes$prop.PA.TOT}
if(prot=="I.IV"){Routes$PA<-Routes$prop.PA.I.IV}


### Run species models
Routes$mainoccG<-factor(Routes$mainoccG, c("For", "Shrub", "Herb"))

ND<-expand.grid(PA=c(0,1), mainoccG=levels(droplevels(Routes$mainoccG))) # Table used for model predicts

NA->effects$For0->effects$For1->effects$Shrub0->effects$Shrub1->effects$Herb0->effects$Herb1


for(spc in 1:(ncol(obsRed)-2)){
  Routes$AB<-obsRed[,spc]
  
  tryCatch({mod<-gam(AB ~ PA * mainoccG + s(Alt_mean, Latitude, Longitude, NPP_mean), family="nb", data=Routes)
  
  NDs<-ND
  NDs$Latitude<-mean(Routes$Latitude[Routes$AB>0], na.rm=T) ; NDs$Longitude<-mean(Routes$Longitude[Routes$AB>0], na.rm=T) ; NDs$Alt_mean<-median(Routes$Alt_mean[Routes$AB>0], na.rm=T) ; NDs$NPP_mean<-median(Routes$NPP_mean[Routes$AB>0], na.rm=T)
  
  if(table(Routes$mainoccG[Routes$AB>0])["Shrub"]> 0){NDs[,7:8]<-predict(mod, newdata=NDs, se.fit=T)} 
  if(table(Routes$mainoccG[Routes$AB>0])["Shrub"]==0){NDs[,7:8]<-NA ; colnames(NDs)[7:8]<-c("fit", "se.fit") ; NDs[c(1,2,5,6),7:8]<-predict(mod, newdata=NDs[c(1,2,5,6),], se.fit=T)} 
  
  effects$For0[spc]<-NDs$fit[NDs$mainoccG=="For"][1] ; effects$For1[spc]<-NDs$fit[NDs$mainoccG=="For"][2]
  tryCatch({effects$Shrub0[spc]<-NDs$fit[NDs$mainoccG=="Shrub"][1] ; effects$Shrub1[spc]<-NDs$fit[NDs$mainoccG=="Shrub"][2]}, error=function(e){cat("Shrubbug")})
  effects$Herb0[spc]<-NDs$fit[NDs$mainoccG=="Herb"][1] ; effects$Herb1[spc]<-NDs$fit[NDs$mainoccG=="Herb"][2]
  
  cat(spc, " ") 
  }, error=function(e){cat("BUG")})
}

# Add sampling variables
effects$N<-v[match(paste0("X",effects$AOU), colnames(obs))]

ech<-data.frame(AOU=effects$AOU)
NA->ech$Ker.For->ech$Ker.Shrub->ech$Ker.Herb # Nb of routes inside the kernell
NA->ech$Pres.For->ech$Pres.Shrub->ech$Pres.Herb # Nb of routes with presence inside the kernell
NA->ech$Ker.PA.For->ech$Ker.PA.Shrub->ech$Ker.PA.Herb # Nb of protected routes inside the kernell
NA->ech$Pres.PA.For->ech$Pres.PA.Shrub->ech$Pres.PA.Herb # Nb of protected routes with presence inside the kernell

for(spc in 1:(nrow(ech))){
  ech[spc, 2:4]<-table(Routes$mainoccG[is.na(obsRed[,spc])==F])[c("For", "Shrub", "Herb")]
  ech[spc, 5:7]<-table(Routes$mainoccG[obsRed[,spc]>0])[c("For", "Shrub", "Herb")]
  try({ech[spc, 8:10]<-table(Routes$prop.PA.TOT[is.na(obsRed[,spc])==F]>0.5, Routes$mainoccG[is.na(obsRed[,spc])==F])["TRUE",c("For","Shrub","Herb")]})
  try({ech[spc, 11:13]<-table(Routes$prop.PA.TOT[obsRed[,spc]>0]>0.5, Routes$mainoccG[obsRed[,spc]>0])["TRUE",c("For","Shrub","Herb")]})
}  

effects$Pres.PA.For<-ech[match(effects$AOU, ech$AOU), "Pres.PA.For"]
effects$Ker.PA.For<-ech[match(effects$AOU, ech$AOU), "Ker.PA.For"]
effects$Pres.For<-ech[match(effects$AOU, ech$AOU), "Pres.For"]
effects$Ker.For<-ech[match(effects$AOU, ech$AOU), "Ker.For"]


write.csv(effects, paste0("effects.species.Protection", prot, ".Species", espe, ".csv"), row.names=F)


### Fig.1 (sample)
effects$PA.For<-(effects$For1-effects$For0)
effects$PA.Shrub<-(effects$Shrub1-effects$Shrub0)
effects$PA.Herb<-(effects$Herb1-effects$Herb0)

png(file = "Plot.sampling.png", width = 20,  height = 13, units = "cm", res = 800)
C=0.75
plot(sign(effects$PA.For)*log(abs(effects$PA.For)) ~ ech$Ker.PA.For, cex=C, col="springgreen4", pch=20, xlab="PAs sampling quality", ylab="PAE", yaxt="n", main="", ylim=c(-log(129000), log(2100))) ; abline(h=0, col="gray41", lty=3, lwd=1.5)
points(sign(effects$PA.Shrub)*log(abs(effects$PA.Shrub)) ~ ech$Ker.PA.Shrub, cex=C, col="firebrick3", pch=20)
points(sign(effects$PA.Herb)*log(abs(effects$PA.Herb)) ~ ech$Ker.PA.Herb, cex=C, col="darkgoldenrod1", pch=20)
axis(2, at=c(-log(10000), -log(1000), -log(100), -log(10), 0, log(10), log(100), log(1000)), labels=c(expression(paste("-10" ^ "4")),expression(paste("-10" ^ "3")),expression(paste("-10" ^ "2")),expression(paste("-10" ^ "1")),0,expression(paste("10" ^ "1")),expression(paste("10" ^ "2")),expression(paste("10" ^ "3"))) ,las=2)
legend("bottomright", pch=20, col=c("springgreen4", "firebrick3", "darkgoldenrod1"), legend=c("Forest", "Shrub", "Herbaceous"), title="Vegetation structure", title.col="gray45")
dev.off()


###############################
### GLOBAL MODELS AND PLOTS ###
###############################
library(DescTools)
library(car)


### Change the line depending on what you want to plot
effects<-read.csv("effects.species.ProtectionTOT.SpeciesNat.Round1.csv") # Plots all PAs, native species only
# effects<-read.csv("effects.species.ProtectionI.IV.SpeciesNat.Round1.csv") # Plots strict PAs, native species only
# effects<-read.csv("effects.species.ProtectionTOT.SpeciesNNNat.Round1.csv")  # Plots all PAs, all species

effects$Latin<-species[match(effects$AOU, species$AOU), "Name"]
effects$PA.For<-(effects$For1-effects$For0)
effects$PA.Shrub<-(effects$Shrub1-effects$Shrub0)
effects$PA.Herb<-(effects$Herb1-effects$Herb0)


### Calculate human-affinity
# Charge obs with all land use types (not only forest, shrub and herbaceous)
obs.full<-read.csv("Obs.NNNative.BBS.Round1.11.01.19.csv", row.names="X") ; obs.full<-obs.full[order(rownames(obs.full)),]
Routes.full<-read.csv("Routes.BBS.Round1.11.01.19.csv") ; Routes.full<-Routes.full[order(Routes.full$RtCode),]
table(rownames(obs.full)==Routes.full$RtCode) # Check 100 % TRUE

obs.full<-obs.full[,names(obs.full) %in% paste0("X",effects$AOU)] ; table(names(obs.full)==paste0("X",effects$AOU))
effects$med.pond.hft<-NA
for(spc in 1:nrow(effects)){
  v<-0
  Routes.full$AB<-obs.full[,spc]
  for(i in 1:nrow(Routes.full)){
    if(replace(Routes.full$AB[i], is.na(Routes.full$AB[i])==T,0)>0){v<-c(v, rep(Routes.full$hft[i], Routes.full$AB[i]))}}
  effects$med.pond.hft[spc]<-median(v, na.rm=T)
}

hist(effects$med.pond.hft)


# Trend. Available in Sauer et al. 2017
trends<-read.csv("Trends_BBS.csv", sep=";")
effects$Trend1966<-trends[match(effects$AOU, trends$AOU.Number), "Trend.1966.2015"]
effects$Trend1966<-Winsorize(effects$Trend1966, probs=c(0.025,0.975), na.rm=T)
effects<-subset(effects, is.na(effects$Trend1966)==F)


### Linear model
M.linear<-lm(PA.For ~ J.main.habitat + Trend1966 + med.pond.hft -1, data=effects)
#M.linear<-lm(Winsorize(PA.Shrub, na.rm=T) ~ J.main.habitat + Trend1966 + med.pond.hft -1, data=effects)
#M.linear<-lm(Winsorize(PA.Herb, na.rm=T) ~ J.main.habitat + Trend1966 + med.pond.hft -1, data=effects)
summary(M.linear)
Anova(M.linear)


### Phylogenetic models
library(phylolm)
library(treeman)

# Subset Jetz consensus tree
birds<-read.tree("Tree.JetzConsensus.15.02.18")
arbre<-as(birds, 'phylo')
arbre$tip.label<-sub("_"," ", arbre$tip.label)
vecteur<-arbre$tip.label
source("D:/Change.Names.Species.R") # Apply name changes due to mismatch of names between Jetz and BBS
arbre$tip.label<-vecteur
Exclude<-arbre$tip.label
Exclude<-subset(Exclude, Exclude %not in% effects$Latin)
arbresub<-drop.tip(arbre, Exclude)
arbresub$tip.label<-sub("_"," ", arbresub$tip.label)


# Exclude species that are not in the analysis
Exclude<-subset(arbresub$tip.label, arbresub$tip.label %not in% droplevels(as.factor(effects$Latin)))
arbresub<-drop.tip(arbresub, Exclude)
effects<-subset(effects, effects$Latin %in% arbresub$tip.label)
rownames(effects)<-effects$Latin

# Create a specific tree for shrub and herb because they have some NAs (the model could not estimate the effect of PAs in herbaceous when there were no protected shrubland in species kernel)
effects.For<-subset(effects, is.na(effects$PA.For)==F)
effects.Shrub<-subset(effects, is.na(effects$PA.Shrub)==F)
effects.Shrub$J.main.habitat<-droplevels(effects.Shrub$J.main.habitat)
effects.Herb<-subset(effects, is.na(effects$PA.Herb)==F)
Exclude.For<-subset(arbresub$tip.label, arbresub$tip.label %not in% droplevels(as.factor(effects.For$Latin)))
Exclude.Shrub<-subset(arbresub$tip.label, arbresub$tip.label %not in% droplevels(as.factor(effects.Shrub$Latin)))
Exclude.Herb<-subset(arbresub$tip.label, arbresub$tip.label %not in% droplevels(as.factor(effects.Herb$Latin)))
arbresub.For<-drop.tip(arbresub, Exclude.For)
arbresub.Shrub<-drop.tip(arbresub, Exclude.Shrub)
arbresub.Herb<-drop.tip(arbresub, Exclude.Herb)

M.phylo.BM<-phylolm(PA.For ~ J.main.habitat + Trend1966 + med.pond.hft -1, data=effects.For, phy=arbresub.For, model="BM", boot=100)
#M.phylo.BM<-phylolm(Winsorize(PA.Shrub, na.rm=T) ~ J.main.habitat + Trend1966 + med.pond.hft -1, data=effects.Shrub, phy=arbresub.Shrub, model="BM", boot=100)
#M.phylo.BM<-phylolm(Winsorize(PA.Herb, na.rm=T) ~ J.main.habitat + Trend1966 + med.pond.hft -1, data=effects.Herb, phy=arbresub.Herb, model="BM", boot=100)
summary(M.phylo.BM)


### Fig.2
Hab<-data.frame(table(effects$J.main.habitat)) ; colnames(Hab)=c("habitat", "N")
Hab$L.mean<-coef(M.linear)[match(Hab$habitat, substr(names(coef(M.linear)),15,100))]
Hab$L.sd<-summary(M.linear)$coefficients[match(Hab$habitat, substr(names(summary(M.linear)$coefficients[,2]),15,100)), 2]
Hab$BM.mean<-coef(M.phylo.BM)[match(Hab$habitat, substr(names(coef(M.phylo.BM)),15,100))]

# IC et ajouts valeurs moyennes
Moy<-c(0,0) 
Hab$L.inf<-Hab$L.mean-1.96*Hab$L.sd ; Hab$L.sup<-Hab$L.mean+1.96*Hab$L.sd 
Hab[,c(3,6,7)]<-Hab[,c(3,6,7)] + Moy[1]*M.linear$coefficients["Trend1966"] + Moy[2]*M.linear$coefficients["med.pond.hft"]    
# Add bootstrap CI for phylolm
Hab$BM.inf<- summary(M.phylo.BM)$coefficients[match(Hab$habitat, substr(names(summary(M.phylo.BM)$coefficients[,2]),15,100)), 4]
Hab$BM.sup<- summary(M.phylo.BM)$coefficients[match(Hab$habitat, substr(names(summary(M.phylo.BM)$coefficients[,2]),15,100)), 5]
Hab[,c(5,8,9)]<-Hab[,c(5,8,9)] + Moy[1]*M.phylo.BM$coefficients["Trend1966"] + Moy[2]*M.phylo.BM$coefficients["med.pond.hft"]    

# graphic setup
par(mar=c(8,4,2,2))
ec=0.15 # Jitter of points
# Set points order
Hab1<-subset(Hab, Hab$habitat %in% c("mixed_forest", "forest", "deciduous_forest", "conifer_forest")) ; Hab2<-subset(Hab, Hab$habitat %not in% droplevels(Hab1$habitat))
Hab<-rbind(Hab1[order(Hab1$BM.mean, decreasing=T),], Hab2[order(Hab2$BM.mean, decreasing=T),])
Hab$habitat<-revalue(Hab$habitat, c("mixed_forest"="Mixed forest", "forest"="Forest", "deciduous_forest"= "Deciduous forest", "conifer_forest"="Conifer forest", "semi_open"="Semi open", "riparian_wetlands"="Riparian", "generalist"="Generalist", "shrub"="Shrub", "arid"="Arid", "open"="Open", "urban"="Urban"))
T=Hab$N/8 # Points size

png(file = "Plot.Habitat.png", width = 20,  height = 13, units = "cm", res = 800)
par(mar=c(7,4,2,2))
plot((1:nrow(Hab))-ec, Hab$L.mean , xaxt="n", xlab="", pch=20, ylim=c(min(Hab$BM.inf)*1.1, max(Hab$BM.sup)*1.1), ylab="Forest PAs effect", col="deepskyblue3", cex=T)
abline(v=4.5, lty=2, col="forestgreen", lwd=2)
points((1:nrow(Hab))-ec, Hab$L.mean, pch=20, col="deepskyblue3", cex=T)
segments((1:nrow(Hab))-ec, Hab$L.inf, (1:nrow(Hab))-ec, Hab$L.sup, col="deepskyblue3")
abline(h=0, col="gray41", lty=3)
axis(1, Hab$habitat, at=1:nrow(Hab), las=2, cex.axis=0.85)
points((1:nrow(Hab)), Hab$BM.mean , pch=20, col="darkorange2", cex=T)
segments((1:nrow(Hab)), Hab$BM.inf, (1:nrow(Hab)), Hab$BM.sup, col="darkorange2")
dev.off()


### Fig.3: Plot Human-affinity
# Plot med.pond.hft 
png(file = "Plot.affinity.png", width = 17,  height = 13, units = "cm", res = 1500)
effects$For.Hab<-effects$J.main.habitat %in% c("forest", "conifer_forest", "deciduous_forest", "mixed_forest")
effects$For.Hab<-replace(effects$For.Hab, as.character(effects$For.Hab)=="TRUE", "forestgreen")
effects$For.Hab<-replace(effects$For.Hab, as.character(effects$For.Hab)=="FALSE", "darkgoldenrod3")
plot(effects$PA.For ~ effects$med.pond.hft, xlab="human-affinity", ylab="Forest PAs effect", pch=20, col=as.character(effects$For.Hab))
abline(h=0, col="gray41", lty=3)
legend("topright", legend=c("Forest species", "Non-forest species"), pch=20, col=c("forestgreen", "darkgoldenrod3"))

NDhft<-data.frame(med.pond.hft=seq(min(effects$med.pond.hft), max(effects$med.pond.hft), length.out=100), Trend1966=0, J.main.habitat="deciduous_forest")
NDhft[,4:5]<-predict(M.linear, type="response", se.fit=T, newdata=NDhft)
NDhft$min<-NDhft$fit-1.96*NDhft$se.fit ; NDhft$max<-NDhft$fit+1.96*NDhft$se.fit
NDhft2<-data.frame(med.pond.hft=seq(min(effects$med.pond.hft), max(effects$med.pond.hft), length.out=100), Trend1966=0, J.main.habitat="semi_open")
NDhft2[,4:5]<-predict(M.linear, type="response", se.fit=T, newdata=NDhft2)
NDhft2$min<-NDhft2$fit-1.96*NDhft2$se.fit ; NDhft2$max<-NDhft2$fit+1.96*NDhft2$se.fit
lines(NDhft$med.pond.hft, NDhft$fit, col="forestgreen", lwd=3)
lines(NDhft$med.pond.hft, NDhft$min, col="forestgreen", lwd=1.5, lty=2)
lines(NDhft$med.pond.hft, NDhft$max, col="forestgreen", lwd=1.5, lty=2)

lines(NDhft2$med.pond.hft, NDhft2$fit, col="darkgoldenrod3", lwd=3)
lines(NDhft2$med.pond.hft, NDhft2$min, col="darkgoldenrod3", lwd=1.5, lty=2)
lines(NDhft2$med.pond.hft, NDhft2$max, col="darkgoldenrod3", lwd=1.5, lty=2)

dev.off()
```
